# Greatwall Kinase regulates Acute Myeloid Leukaemia Cell Division through a Non-Canonical Mechanism

**DOI:** 10.64898/2026.03.16.712046

**Authors:** Sandra M. Martin-Guerrero, Tommy Shields, Pedro Casado, Róbert Zach, Vinothini Rajeeve, Nadia Afroz-Nishat, Irbaz I. Badshah, Megan Meredith, William R Foster, Helfrid Hochegger, Pedro R. Cutillas

## Abstract

Greatwall kinase regulates mitotic progression by phosphorylating ENSA and ARPP19, thereby inhibiting PP2A-B55. Moreover, Greatwall has been implicated in oncogenesis, particularly in solid tumours, but the mechanisms by which Greatwall regulates the cell cycle in other malignancies remain unclear. Here, we show that Greatwall regulates cytokinesis and cell cycle progression in acute myeloid leukaemia (AML) cells through a pathway distinct from ENSA-PP2A-B55. AML cells require Greatwall expression and activity to proliferate, as revealed by pharmacological and systematic genetic perturbation experiments. Mechanistically, Greatwall inactivation or genetic depletion does not measurably affect the ENSA–PP2A-B55 pathway. Instead, loss of Greatwall function alters cytokinesis, and the phosphorylation of proteins involved in cytoskeletal organisation and cytokinesis, including MARK3, which we identify as a direct Greatwall substrate in AML cells. Together, these findings reveal that the Greatwall kinase signalling network is wired differently in leukemic cells, thus uncovering a novel of cell cycle regulation.

## INTRODUCTION

Greatwall (Gwl), also known as MASTL (microtubule-associated serine/threonine kinase-like), is a master regulator of cell cycle progression (Cundell et al, 2013). Activated at the G2/M transition, Gwl phosphorylates ENSA (α-endosulfine) and ARPP19 (cAMP-regulated phosphoprotein 19) at Ser62 and Ser67, respectively (Cundell et al, 2013; Alvarez-Fernandez et al, 2013; Hégarat et al, 2014; Vigneron et al, 2009). Phosphorylated ENSA and ARPP19 bind to the PP2A-B55 phosphatase complex and inhibit its activity, thereby preventing premature dephosphorylation of mitotic substrates (Castro & Lorca, 2018; Cundell et al, 2013). This inhibition enables CDK1-cyclin B to establish and maintain the high-phosphorylation state required for mitotic entry and progression (Goldberg, 2010; Rata et al, 2018; Williams et al, 2014; Vigneron et al, 2009). Consistent with this central role in mitotic control, Gwl is frequently deregulated in cancer, and its inhibition disrupts cancer cell proliferation and survival, highlighting it as a potential therapeutic target (Burgess et al, 2010; Castro & Lorca, 2018; Misra et al, 2023; Peng et al, 2010; Sanz-Castillo et al, 2023; Uppada et al, 2018; Vera et al, 2015; Wong et al, 2016; Zach et al, 2025). However, its role in haematological malignancies such as acute myeloid leukaemia (AML) remains poorly characterised, despite evidence from large-scale functional screens identifying Gwl as essential for AML survival (Tzelepis et al, 2016).

AML is characterised by impaired differentiation and clonal expansion of myeloid progenitor cells, ultimately leading to bone marrow failure (Khwaja et al, 2016). There were an estimated 144,645 new AML cases worldwide in 2021 (Zhou et al, 2024). Advances in our understanding of AML biology have led to the gradual introduction of new therapeutic approaches, including targeted agents directed against key disease drivers such as FLT3, BCL2, and IDH1/2, for molecularly defined patient subgroups (Döhner et al, 2022). However, despite these advances, clinical responses remain variable and overall survival is still poor (Casado & Cutillas, 2023; Zhou et al, 2024). This highlights an unmet need for therapies that target additional vulnerabilities in AML biology. Although cell cycle regulatory mechanisms have not yet been widely exploited therapeutically in AML, defining AML-specific dependencies in cell cycle control may uncover novel treatment opportunities and improve our understanding of how cell cycle dysregulation contributes to leukaemogenesis.

In this study, we took advantage of a novel small-molecule inhibitor (namely, c604 (Zach et al, 2025)) with target selectivity for Gwl to define the function of this kinase across 14 AML cell lines and eight primary AML cultures directly taken from patients. Key findings were validated using shRNA-mediated Gwl knockdown. Inhibition of Gwl markedly impaired leukemic cell proliferation and mitotic progression, leading to cytokinesis defects including polyploidy and multinucleation. Notably, these effects occurred independently of ENSA/ARPP19 (hereafter referred to as ENSA) and PP2A-B55. In contrast to mechanisms observed in solid cancer cells, upon Gwl inhibition, AML cells showed no changes in ENSA phosphorylation or PP2A-B55 association across the cell cycle or following pharmacologic or genetic perturbation of Gwl. Instead, we identify a non-canonical mechanism by which Gwl regulates cytokinesis through direct phosphorylation of proteins involved in cytokinesis. Of these, we confirmed that MARK3 activity is altered upon Gwl inhibition. These findings reveal distinct wiring of Gwl signalling in AML and highlight a key mechanism of cell cycle control in leukaemia cells.

## RESULTS

### Gwl activity is required for AML cell viability and proliferation

To assess the anti-leukemic effects of Gwl inhibition in AML, we treated 14 different AML cell lines with c604, a selective Gwl inhibitor previously characterised in solid tumour cells (Zach et al, 2025). Gwl inhibition reduced the proliferation of all tested AML cell lines, except for HL-60 cells (Fig. EV1A, B). However, we observed a heterogeneous response in the reduction of cell viability across these models (Fig. 1A), with MOLM-13 and MV4-11 (FLT3-ITD mutant-positive) being particularly sensitive to c604, whereas KG-1 and HL-60 cells were relatively resistant (Fig. 1A, B). The anti-leukaemic impact of Gwl in AML was confirmed with data from systematic genetic targeting of this kinase across cancer cell lines (Fig. 1C, Fig. EV1C), showing that AML cells exhibit high genetic dependency to Gwl relative to other tumour types (Fig. 1C and Fig. EV1C, and (Tzelepis et al, 2016)).

**Figure 1.**
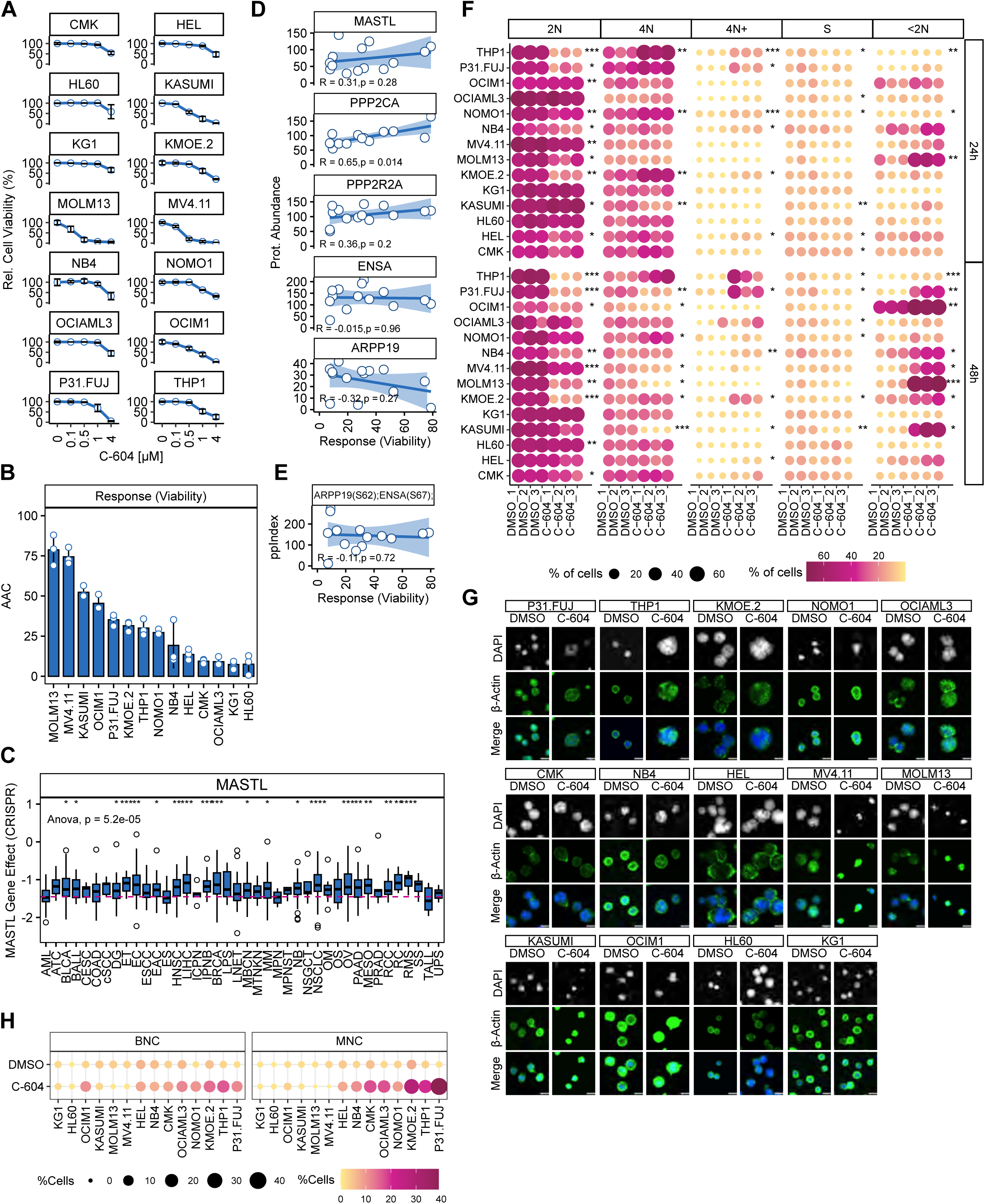
Reduction in Gwl activity decreases cell viability and alters cell cycle progression in AML cells. (A) Cell viability of AML cell lines treated with different concentrations of the Gwl inhibitor, compound c604, for 72 hours, relative to vehicle-treated cells. Values represent mean ± SD of 3-4 biological replicates. (B) Response to c604 calculated as Area Above the Curve (AAC) from the data presented in A. (C) Gwl genetic dependency of cancer cell lines grouped by primary disease from CRISPR Chronos Dataset. Statistical significance was calculated by unpaired t-Test between AML versus the rest of the cancer types (\**: p<0.05, **: p<0.01 and **\*\*\****: p<0.001). (D) and (E) Relationship between the response calculated for c604 and the protein levels of the named proteins or phosphorylation sites. rho coefficient and statistical significance (by Spearman) are shown. (F) Distributions of cell cycle groups (2N, S, 4N, 4N+ and <2N) in AML cells treated with c604 (2 µM) for 24 and 48 hours of 3 independent biological replicates. Statistical significance was calculated by unpaired t-Test between treated cells versus non-treated (DMSO) groups (\**: p<0.05, **: p<0.01 and \*\*\** : p<0.001). (G) Representative Spinning Disk confocal microscopy images of AML cells after 48 hours of c604 treatment (2 µM). Nuclei were stained with DAPI (4’,6-diamidino-2-phenylindole, shown in grey scale upper panels) and cells immunostained for β-actin (green). Scale bar: 10 μm. (H) The presence of multiple nuclei was analysed after 48 hours of c604. Data points show the percentage of binucleated cells (BNC) or multinucleate cells (MNC) over the total of cells analysed for each condition after 48 hours of treatment with c604 or DMSO (n=260-1919 cells).

### PPP2CA expression is linked to responses of AML cells to c604

A recent study linked c604 sensitivity to low Gwl and high B55α expression in solid tumour cell lines (Zach et al, 2025). Contrary to these findings, we observed no significant correlation between c604 sensitivity and Gwl or B55α protein levels, or phosphorylation of Gwl substrates in AML cells (Fig. 1D, E and Fig. EV1D). In contrast, the antiproliferative effects of c604 showed a significant correlation with protein levels of the PP2A catalytic subunit (PPP2CA; Fig. 1D). Thus, our data suggest that Gwl/PPP2CA expression may represent a candidate biomarker for stratifying anti-Gwl treatments in AML

### Gwl inhibition induces cell death and polyploidisation in AML cell lines

Gwl inhibition disrupts cell cycle progression in solid tumours (Bisteau et al, 2020; Vigneron et al, 2009; Zach et al, 2025; Alvarez-Fernandez et al, 2013). We therefore assessed cell cycle progression in 14 AML cell lines after 24 and 48 hours of c604 treatment (2 µM; Fig. 1F). Interestingly, in the most sensitive AML cell lines (MV4-11, MOLM-13, Kasumi-1, OCI-M1), Gwl inhibition did not induce accumulation of 4N DNA content; instead, SubG1 events (<2N) predominated, indicating high cell death (Fig. 1F). Lower inhibitor doses (0.25 µM) yielded similar results (Fig. EV1E). In contrast, several other AML lines (P31/Fuj, KMOE-2, THP-1, NOMO-1, and to a lesser extent CMK, HEL, NB4) displayed increased 4N DNA content after 24- or 48-hour treatment (Fig. 1F), consistent with the impact of the compound on cell viability (Fig. 1A). c604 treatment induced binucleated (BNC) or multinucleated (MNC) cells in most AML models (Fig. 1G, H). Intriguingly, cells with a FLT3-ITD genotype (MV4-11 and MOLM-13 cells) and Kasumi-1 cells did not exhibit this phenotype (Fig. 1G, H), showing instead a clear reduction in cell size and chromatin condensation, which is indicative of cell death.

To investigate the differential responses to c604 in greater detail, we used LC-MS/MS phosphoproteomics (Zach et al, 2025; Gerdes et al, 2021; Casado et al, 2023) to profile the impact of 2 µM compound treatment on asynchronous cells. We focused on models observed to be resistant (HL-60), sensitive with multinucleation (P31/Fuj), or sensitive without multinucleation (MV4-11) to c604 (Fig. 2A). We chose a 2-hour time point to capture signalling changes while minimising cell death due to Gwl inhibition. Notably, the impact of c604 on the phosphoproteome reflected the phenotypic responses to the compound: HL-60 displayed the fewest changes in protein phosphorylation, while MV4-11 and P31/Fuj showed a decrease or increase in protein phosphorylation, respectively (Fig. 2B).

**Figure 2.**
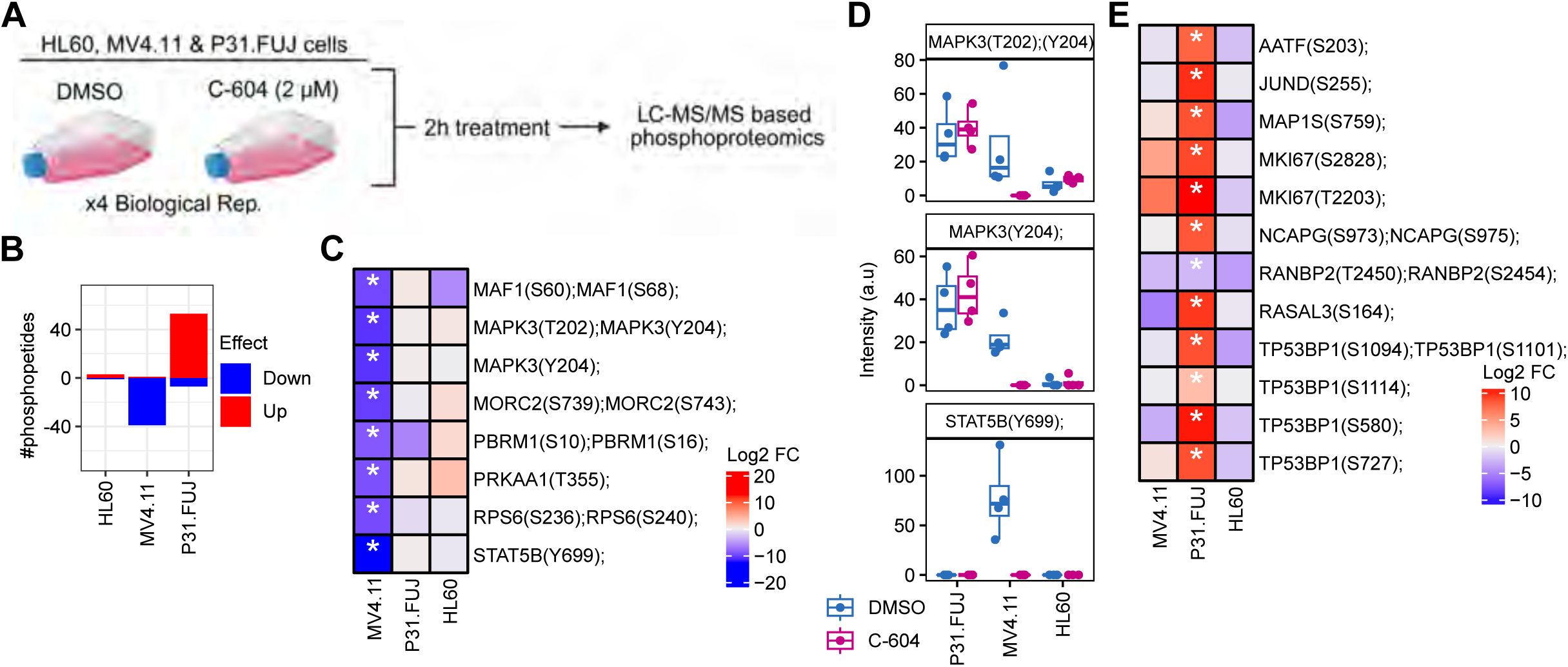
c604 impacts protein phosphorylation in asynchronous AML cells in a context-dependent manner. (A) Asynchronous AML cells (MV4-11, HL-60 and P31/Fuj cells) were treated with c604 (2 µM) or DMSO for 2 hours and then processed for LC-MS/MS based phosphoproteomic analysis (n=4 independent experiments). (B) Number of significant phosphopeptides identified for each cell line compared to DMSO-treated cells (Log2 FC > |0.8|, p<0.05 & FDR<0.15). (C) Representative phosphopeptides significantly modulated by the treatment in MV4-11 cells linked to FLT3-ITD signalling. (D) Normalised signal intensity of the named STAT5 and ERK1 (MAPK3) phosphosites from C. (E) Representative phosphopeptides significantly modulated by c604 treatment in P31/Fuj cells.

In MV4-11 cells (FLT3-ITD mutant-positive), c604 decreased the phosphorylation of STAT5 signalling proteins, including STAT5B at Tyr699 (Fig. 2C), at the activation sites of ERK2 (MAPK3 Thr202/Tyr204) and other proteins downstream of FLT3 and STAT5 signalling (Fig. 2C). c604 did not reduce STAT5 or ERK2 phosphorylation in FLT3-ITD-negative cells (P31/Fuj and HL-60; Fig. 2C, D), suggesting disruption of FLT3 signalling in FLT3-ITD-mutant cells. In AML, FLT3-ITD mutations activate oncogenic STAT5 signalling essential for leukemic proliferation and survival (Choudhary et al, 2007; Spiekermann et al, 2003; Nelson et al, 2012; Vainchenker & Constantinescu, 2013; Wang & Bunting, 2013; Wierenga et al, 2006). This may explain the sensitivity of MOLM-13 and MV4-11 cells to c604, although an off-target effect on FLT3-ITD cannot be excluded.

In FLT3-ITD-negative P31/Fuj cells, c604 increased phosphorylation of proteins involved in mitosis, DNA repair, and G2/M checkpoint (Fig. 2E), including cohesion complex components (NCAPG), MKI67, microtubule-associated proteins (MAP1S), and DNA damage response factors (TP53BP1; Fig. 2B). These changes were consistent with 4N DNA accumulation and defective mitotic progression. Overall, these results show that pharmacological Gwl inhibition induces mitotic arrest in AML cells via context-dependent mechanisms that, depending on the AML subtype, involve a decrease in STAT5 signalling or a disruption of the mitotic machinery.

### ENSA phosphorylation is unaffected by Gwl inhibition in AML

c604 alters solid tumour cells’ survival and cell cycle progression through the well-established ENSA-PP2A-B55 axis (Zach et al, 2025). To assess the impact of Gwl inhibition on leukaemic mitosis, we performed phosphoproteomic analysis of prometaphase-arrested AML cells and HeLa cells treated with c604 (Fig. 3A). To promote mitotic exit, AML cells were treated with CDK1 inhibitor RO-3306 (10 µM, 2 hours), as previously described (Hegarat et al, 2014). Unsynchronised P31/Fuj cells treated with DMSO were included as controls. Of the 17,328 phosphopeptides detected, mitotic arrest induced the phosphorylation of 3,697 peptides in P31/Fuj cells, whereas CDK1i reduced the phosphorylation of 2,729 peptides (p<0.05, FDR<0.15, Fig. 3B). c604 also decreased the phosphorylation of 455 peptides in P31/Fuj cells (p<0.05, FDR<0.15, Fig. 3B). As previously described (Zach et al, 2025), c604 reduced the phosphorylation of 1,096 peptides in HeLa cells (p<0.05, FDR<0.15; Fig. 3B), thus serving as a positive control for the experiment.

**Figure 3.**
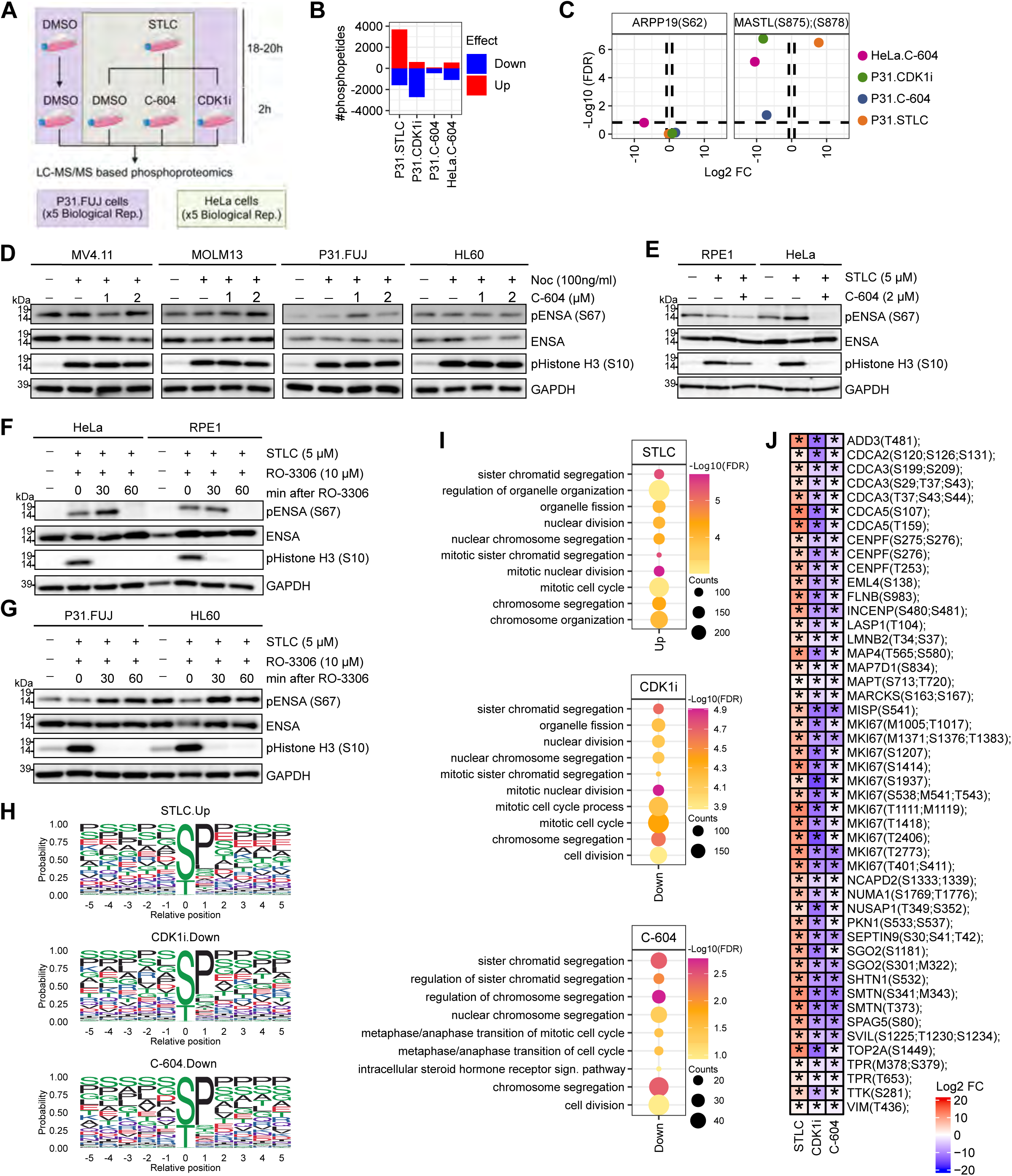
Phosphoproteomics of c604 treated mitotic cells reveals AML-specific mechanisms of cell cycle regulation. (A) P31/Fuj cells, arrested in mitosis with STLC (5 µM) for 18-20 hours, were treated with c604 (2 µM), CDK1 inhibitor, RO-3306 (10 µM) or DMSO and collected for phosphoproteomic analysis (n=5 independent experiments). Asynchronous P31/Fuj treated with DMSO were also included. HeLa cells arrested in mitosis were included as positive controls. Mitotic cells were collected by mitotic shake off, treated with c604 (2 µM) for 2 hours and collected for phosphoproteomic analysis (n=5). (B) Number of phosphopeptides significantly modulated by each condition (Log2 FC > |0.8|, p<0.05 & FDR<0.15). (C) FC (expressed in Log2) and FDR values (-Log10 FDR) of ARPP19 (Ser62) and Gwl (Ser875, Ser878) phosphopeptides in the indicated cell lines and treatments. (D-E) AML cells, RPE1 and HeLa cells, arrested in mitosis using nocodazole (100 ng/ml) or STLC (5 µM) for 18-20 hours, were treated with c604 (1-2 µM) for 2 hours and processed for western blot analysis. Representative western blot images are shown. (F-G) P31/Fuj, HL-60, RPE1 and HeLa cells, arrested in mitosis as before, were treated with CDK1 inhibitor, RO-3306 (10 µM) for 0, 30 and 60 min and collected for western blot analysis. Representative western blot images are shown. In D, E, F and G, ENSA and pENSA refer to total and phosphorylated ENSA/ARPP19 levels, respectively. Phosphorylation of Histone 3 at Ser10 (pHistone H3 S10) represents a mitotic marker. GAPDH was used as a loading control. (H) Phosphorylation motifs of identified significantly upregulated (STLC) or downregulated (CDK1i and c604) in P31/Fuj cells. (I) Top10 Gene ontology terms significantly enriched in phosphopeptides significantly upregulated (STLC) or downregulated (CDK1i and c604) in P31/Fuj cells. (J) Representative phosphopeptides significantly downregulated by c604 in P31/Fuj cells.

Mitotic arrest strongly increased Gwl phosphorylation at Ser875/Ser878, whereas CDK1i and c604 markedly decreased it in P31/Fuj and HeLa cells (Fig. 3C), confirming mitotic Gwl activation (Hermida et al, 2020) in both cell types and ruling out pharmacokinetic differences across models. Despite effective Gwl inhibition, ENSA phosphorylation was largely unchanged in AML cells, but not in HeLa cells (p=0.0153, FDR=0.156, Fig. 3C). In P31/Fuj cells, ENSA phosphorylation did not increase at mitosis compared to asynchronous cells (Log2 FC = -0.21, p=0.95, FDR=0.97) and was unaffected by c604 or CDK1i (Fig. 3C). Similar patterns were observed across additional AML lines (Fig. 3D) and contrasted with ENSA phosphorylation in non-haematological cells (HeLa, RPE1), which peaked at G2/M and declined upon CDK1i or c604 treatment (Fig. 3E, F) as previously reported (Zach et al, 2025; Hegarat et al, 2014). In addition, CDK1i did not decrease ENSA phosphorylation across time in AML cells (Fig. 3G). These data reveal an unexpected uncoupling between Gwl activity and ENSA phosphorylation in AML, indicating that ENSA phosphorylation can occur independently of Gwl kinase activity and mitotic phase, highlighting noncanonical mitotic signalling dynamics in leukaemic cells.

In P31/Fuj cells, mitotic arrest increased phosphorylation of CDK1-linked Ser/Thr-Pro (S/T-P) motifs and mitotic proteins (Errico et al, 2010; Songyang et al, 1994), whereas CDK1i reduced it (Fig. 3H, I). c604 decreased phosphorylation of these motifs and proteins, particularly those involved in chromosome segregation, in AML and HeLa cells (Fig. 3H, I, Fig. EV1F). c604 specifically reduced phosphorylation of proteins involved in spindle alignment and chromosome segregation (Liao et al, 1995; Mills et al, 2017; Nakano et al, 2010; Stucke et al, 2002; Sun et al, 2021; Sun & Kaufman, 2018; Wheatley et al, 2001), including TTK/MPS1, TPR, MKI67, NUMA1, NUSAP, INCENP, and CENPF (Fig. 3J). Our results, therefore, show that Gwl inhibition impairs the biochemical composition of proteins involved in microtubule organisation, spindle stability, and cytokinesis, likely leading to mitotic errors, and explaining the observed effects of Gwl inhibition on multinucleation and cell death (Fig. 1).

### Gwl knockdown alters mitotic phosphorylation independently of ENSA in AML

Next, we examined whether genetic reduction of Gwl recapitulates the effects of c604. To this end, we generated inducible knockdown cell lines for Gwl (shGwl) and scrambled shRNA (shCtrl) in P31/Fuj cells (Fig. 4A). We induced the expression of shRNAs for Gwl and control and then subjected the cells to prometaphase arrest (Fig. 4B). Additional controls included cell lines without induction and DMSO-treated cells as asynchronous cells (Fig. 4B). LC-MS/MS phosphoproteomics revealed that mitotic arrest (STLC vs. DMSO) induced the expected increase in phosphorylation in control cell lines (shCtrl = 1869; shCtrl + IPTG = 1802; shGwl = 2008 phosphopeptides, FC>0.8, p<0.05, and FDR<0.15, Fig. 4C). Notably, Gwl knockdown limited mitotic phosphorylation, indicating dependence of these sites on Gwl activity (shGwl + IPTG = 1371 phosphopeptides, Fig. 4C). Specifically, in shGwl cells, 191 phosphosites did not respond to STLC as they did in control cells (Fig. 4D). Gene Ontology analysis revealed that these phosphosites belonged to proteins linked to chromosome segregation and organisation of the nuclear envelope, with the (S/T-P) motif the most represented (Fig. 4E, F). STLC increased phosphorylation of proteins involved in chromosome condensation and segregation (e.g. MKI67, CENPV, NCAPH, and TPR) in shCtrl cells, an effect blunted in shGwl cells (Fig. 4G). A similar pattern was observed for proteins regulating spindle organisation, nuclear architecture, microtubule dynamics, and cytokinesis (Fig. 4H, K). These findings recapitulate those seen with c604 and indicate that reduced Gwl activity disrupts the phosphorylation of proteins required for mitotic progression.

**Figure 4.**
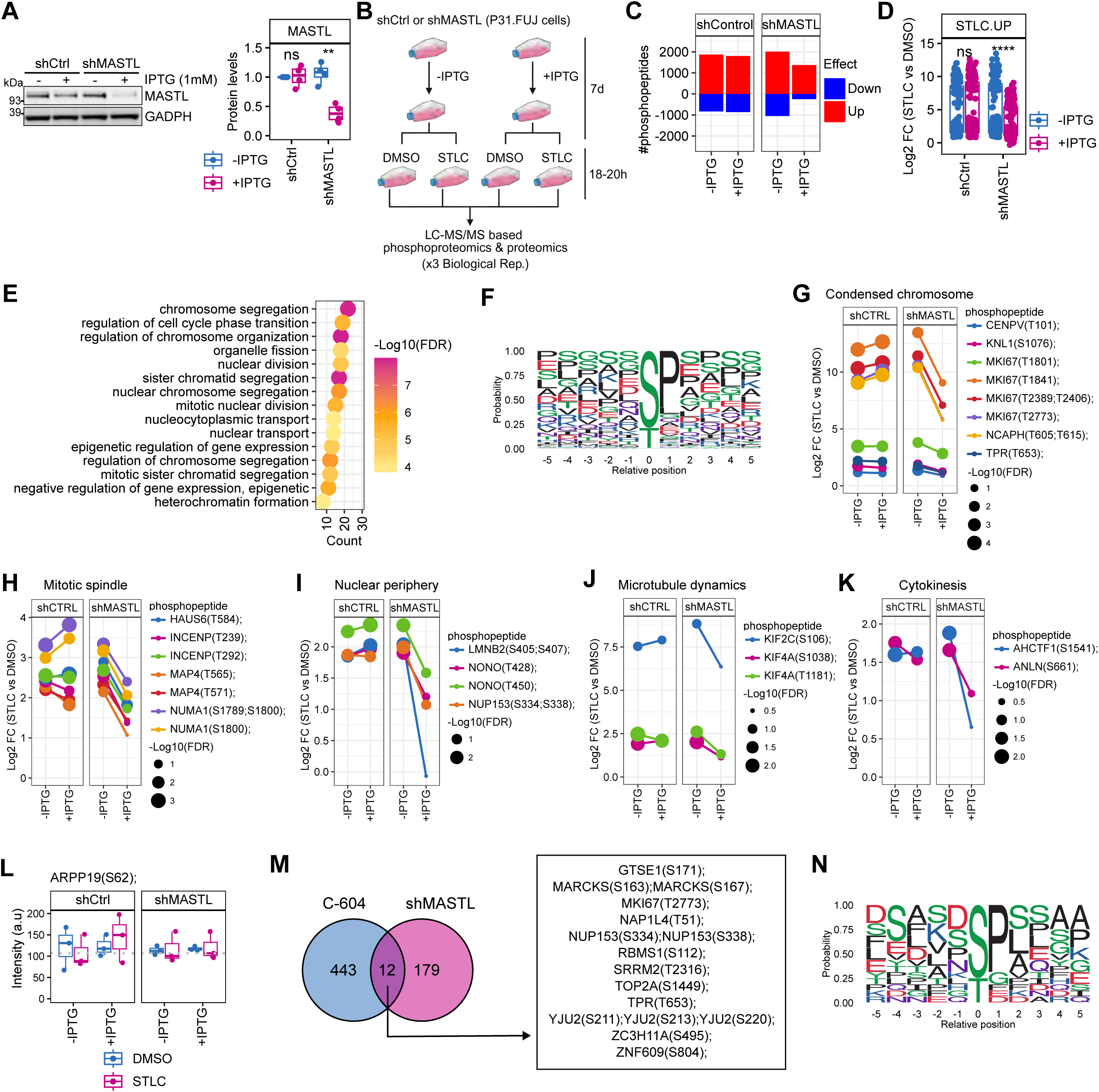
Dissection of AML-specific mitotic regulatory mechanisms downstream of Gwl. (A) WB densitometric analysis of 4 independent biological replicates confirming inducible Gwl knockdown in P31/Fuj cells. shRNA targeting Gwl or scrambled control (shCtrl) was induced by the addition of 1 mM IPTG for 7 days. Assessment of significance was by unpaired t-test (p<0.01). (B) Scheme of treatments of stable cell lines subjected to phosphoproteomic analysis. shCtrl and shGwl cell, previously incubated with 1 mM IPTG for 7 days, were treated with DMSO or STLC 5 µM (mitotic arrest) for 18-20 hours and processed for phosphoproteomics. No induced cells (- IPTG) for shCtrl and shGwl cell lines were included as controls. (C) Number of significant phosphopeptides identified in each condition for P31/Fuj (STLC vs DMSO, Log2 FC > |0.8|, p<0.05 & FDR<0.15). (D) Distribution of the Log2 FC (STLC vs DMSO) of common significant phosphopeptides in shCtrl cells (±IPTG) and shGwl -ITPG that were increased upon STLC treatment. Estimation of statistical significance of differences between induced (+IPTG) versus non-induced (- IPTG) groups was by unpaired t-Test. (E) Top 10 Gene ontology terms significantly enriched the phosphopeptides indicated in D. (F) Sequence motifs of phosphopeptides identified in D. Examples of phosphopeptides linked to chromosome condensation (G), mitotic spindle organisation (H), nuclear periphery (I), microtubule dynamics (J) and cytokinesis (K). Dot plots represent the Log2 FC of each phosphopeptide (STLC vs DMSO) in each cell line. Data point sizes represent the - Log10(FDR) for the STLC vs DMSO comparison. (L) Distribution of the normalised signal intensity for ARPP19 Ser62. (M) Overlap between the significant phosphopeptides identified upon Gwl inhibition (c604) and Gwl knockdown. (N) Sequence motifs of common phosphopeptides between c604 and shGwl.

Consistent with results obtained with c604, ENSA phosphorylation remained unchanged in shGwl cells, both asynchronously and after STLC treatment (Fig. 4L). We also compared the 191 sites affected by Gwl knockdown (Fig. 4D) with those reduced after c604 treatment in mitotic P31/Fuj cells (Fig. 3B). Despite differences in the approaches, we identified 12 concordant phosphosites in proteins involved in nuclear transport or nuclear envelope organisation (e.g., TPR and NUP153), mitotic progression (e.g., MKI67), and microtubule organisation (e.g., MARCKS) (Fig. 4M). In addition, these 12 sites showed phosphorylation mainly within the (S/T-P) motif (Fig. 4N), as observed in the phosphoproteome globally reduced by c604. These results identify a set of proteins downstream of Gwl that may contribute to its anti-mitotic effects.

### Gwl inhibition does not affect ENSA-B55α binding in AML

Canonically, phosphorylated ENSA binds to the regulatory subunit B55 and inhibits the PP2A-B55 complex by “unfair competition” (Cundell et al, 2013; Hached et al, 2019; Mochida, 2014). Therefore, if ENSA phosphorylation is unaltered in AML cells, binding of ENSA to B55 should remain intact. To test this hypothesis, we measured the levels of ENSA bound to B55α in asynchronous AML cells and in mitotic AML cells treated with vehicle, c604, or CDK1i (Fig. 5). As a control, we also evaluated the levels of ENSA bound to B55α in mitotic HeLa cells treated with vehicle or c604 (Fig. 5A). As expected, the binding of ENSA to B55α in mitotic HeLa cells (STLC treatment) was dependent on Gwl activity (Fig. 5A). In contrast, and consistent with our previous results, we observed no significant changes in ENSA binding to B55α in P31/Fuj and KMOE-2 cells upon Gwl inhibition (Fig. 5B, C), which showed binding to B55α in all the conditions tested. These data confirm that, unlike in other systems, Gwl kinase modulates mitotic progression independently of the ENSA-PP2A-B55α axis in AML cells, and support a distinct mechanism through which Gwl kinase regulates mitotic progression in this tumour type.

**Figure 5.**
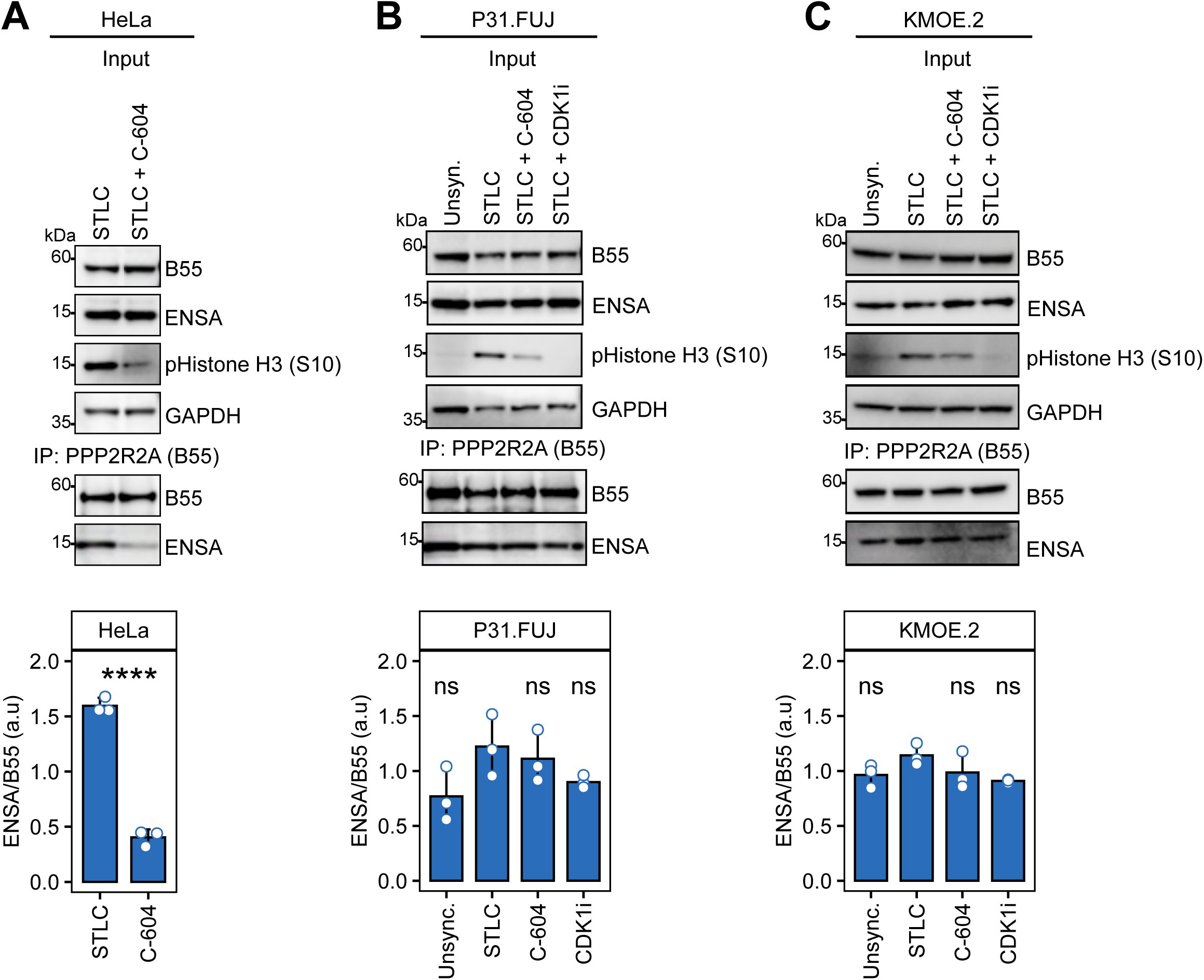
ENSA-B55α physical interaction is cell-cycle-dependent in HeLa cells but not in AML cells. (A) Mitotic HeLa cells (STLC treatment, 5 µM), treated with DMSO or c604 (2 µM) for 2 hours, were subjected to immunoprecipitation assays (IPs) against B55α. Co-immunoprecipitated ENSA was measured by western blot. Representative western blot images and results of 3 independent experiments are shown. Data points represent the ratio of ENSA bound to B55α. (B) Asynchronous P31/Fuj and mitotic arrest P31/Fuj cells (STLC treatment, 5 µM), treated with DMSO, c604 (2 µM) or CDK1i (10 µM) for 2 hours, were subjected to IPs as in A. (C) As in B, but using asynchronous KMOE-2 as a model. An unpaired t-test was used to assess the statistical significance of differences between treatments and the STLC control (***: p<0.001).

### Gwl inhibition does not alter ENSA phosphorylation in primary AML cells

As shown above, Gwl inhibition impaired cell cycle progression in AML cell lines independently of ENSA. We next asked whether this mechanism also operates in primary AML cells (Fig. 6A). Ex-vivo treatment of mononuclear cells directly obtained from AML patients showed that c604 reduced the viability of six of eight cases tested (Fig. 6B, C). Phosphoproteomics revealed that c604 induced changes in protein phosphorylation in each primary AML sample (Fig. 6D). Interestingly, as observed in AML cell lines, c604 did not modify ENSA phosphorylation (Fig. 6E). Although the basal phosphorylation of ENSA varied across patients, this phosphorylation was elevated at basal conditions (Fig. 6E). We observed altered phosphorylation of proteins related to cell cycle, such as NUMA1, MYPT1, NIPBL, and PABIR1, as well as altered phosphorylation of MAP kinase p38 alpha (MAPK14), GSK3β and phosphatase SSH1 (Fig. 6F). These results in primary AML cells reinforce the observations derived from cell line experiments, indicating a decoupling between Gwl activity and ENSA phosphorylation in AML.

**Figure 6.**
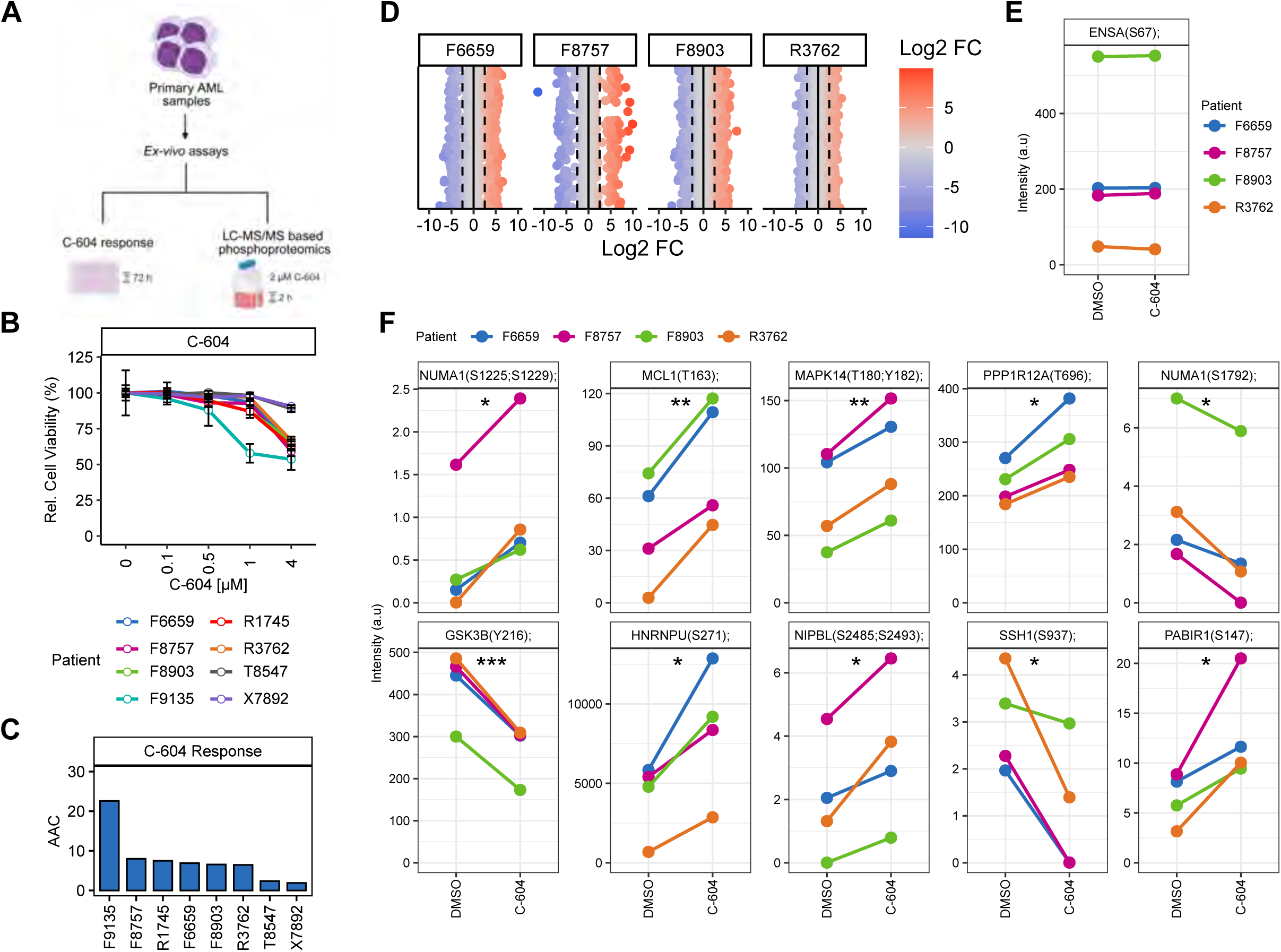
The Gwl inhibitor c604 reduces viability in primary AML cells and broadly alters their phosphoproteome without reducing ENSA phosphorylation. (A) Primary mononuclear cells from eight AML patients were subjected to drug response assays with c604 and four underwent phosphoproteomic analysis. (B) Relative cell viability of AML primary cells treated with different concentrations of compound c604, for 72 hours. Values represent mean ± SD of 3-4 technical replicates for each patient sample. (C) Response to c604 calculated as Area Above the Curve (AAC) from the data presented in B. (D) AML primary samples were treated with 2 µM of c604 or DMSO for 2 hours and processed for LC-MS/MS-based phosphoproteomics. Datapoints represent Log2 FC of phosphopeptides identified. (E) Normalised signal intensity obtained from LC-MS/MS for ENSA phosphorylated at Ser67. (F) Normalised signal intensity obtained from LC-MS/MS for representative phosphopeptides. In E and F, paired t-Test was used to calculate the statistical significance of differences between c604 and DMSO (\**: p<0.05, **: p<0.01 and **\*\*\****: p<0.001).

### Gwl inhibition alters cytoskeletal protein phosphorylation

To investigate how Gwl impairs cell cycle progression in AML, we performed in-cell kinase assays (as previously described (Al-Rawi et al, 2023)) in fixed AML cells to identify noncanonical substrates of this kinase. Fixed and permeabilised cells were incubated with recombinant active Gwl (8xHis-Gwl-2xFlag or GST-Gwl) plus ATP, and the resulting phosphorylation was analysed by LC-MS/MS (Fig. 7A). Cells incubated with buffer alone served as negative controls. Both kinases induced a moderate increase in global serine phosphorylation, with GST-Gwl producing a higher number of Gwl phosphopeptides, consistent with greater kinase activation (Log2 FC >0.7, p<0.05; Fig. 7B, D and Fig. EV2A, B). Interestingly, both kinase assays showed increased phosphorylation of MARK3 (MAP/microtubule affinity-regulating kinase 3; Fig. 7E, Fig. EV2B). Because of its relevance in microtubule dynamics and in cell cycle progression (Drewes et al, 1998; Machino et al, 2022; Ogg et al, 1994; Peng et al, 1998), we sought to validate whether Gwl directly phosphorylates MARK3, as suggested by our results. Remarkably, an in vitro kinase assay using recombinant MARK3 and Gwl (Fig. 7F) revealed that Gwl directly phosphorylates several MARK3 residues (Fig. 7F) involved in 14-3-3 binding (Goransson et al, 2006).

**Figure 7.**
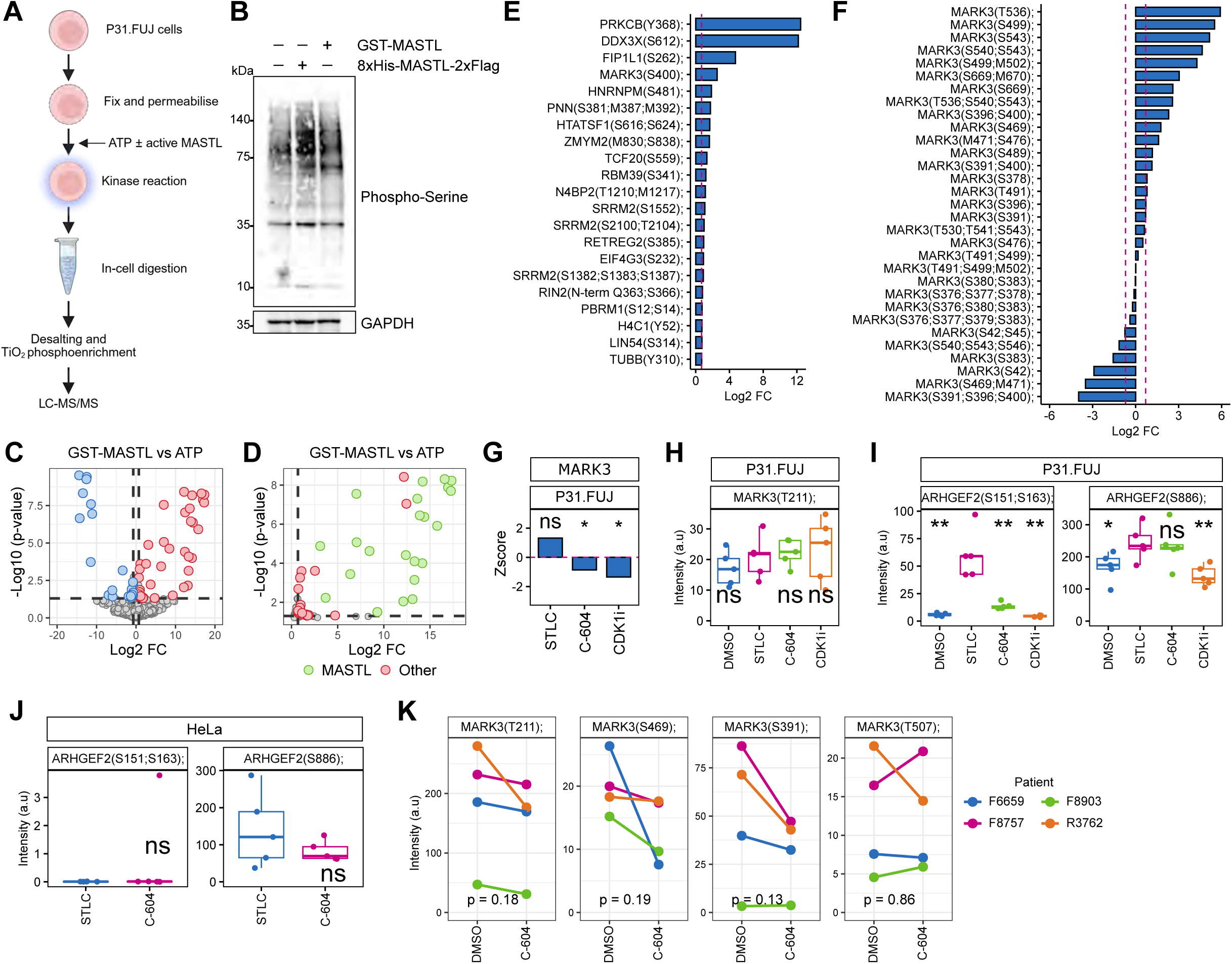
Gwl phosphorylates MARK3 kinase, and c604 decreases MARK3 substrate phosphorylation. (A) Scheme of the in vitro kinase assay on fixed P31/Fuj cells. (B) Western blot image of phosphorylated proteins in serine residues (phosphor-Serine) in P31/Fuj cells incubated only with ATP or recombinant active Gwl kinases (8xHis-Gwl-2xFLAG or GST-Gwl). GAPDH was used as loading control. (C) Phosphopeptides identified by LC-MS/MS in the kinase assay on fixed P31/Fuj cells. X-axis represents Log2 FC between cells incubated with GST-Gwl versus those incubated with only ATP (n=3), with significant phosphopeptides highlighted (p<0.05, Log2 FC > |0.7|). (D) Datapoints of phosphopeptides belonging to Gwl (green) from the volcano plot shown in C. (E) Significant phosphopeptides identified in the kinase assay ranked by Log2 FC. Dashed pink line indicates the Log2 FC threshold of 0.7. (F) In vitro kinase assay using recombinant MARK3 and Gwl kinases was performed and analysed by LC-MS/MS (n=2). Shown are the Log2 FC of MARK3 phosphopeptides. Dashed pink line indicates the Log2 FC threshold of 0.7. (G) Overall MARK3 phosphorylation of sites in the protein regions involved in 14-3-3 binding in P31/Fuj cells was estimated as the Z-score across the conditions. (H) Distributions of the normalised signal intensity for MARK3 phosphorylation at the T-loop activation site (Thr211) in P31/Fuj cells. (I-J) Normalised signal intensity for ARHGEF2 phosphorylation at Ser151 and Ser163 or at Ser886 in P31/Fuj cells (I) and in HeLa cells (J). (K) Normalised signal intensity for MARK3 phosphopeptides identified in AML primary samples treated with c604. Statistical significance of differences between groups was estimated by unpaired t-Test (\**: p<0.05,** : p<0.01 and **\*\*\**** : p<0.001).

To determine whether Gwl inhibition affects MARK3 phosphorylation at 14-3-3 binding sites in mitotic cells, we derived a phosphorylation site enrichment score (Ser396, Ser400, Ser419, Ser469) as a Z-score from our phosphoproteomic dataset (Fig. 7G, Fig. EV2C). Mitotic arrest tended to increase the phosphorylation at these sites, which were reduced by c604 and CDK1i, consistent with kinase assay results (Fig. 7G). Phosphorylation of the activation loop (T-loop, Thr211) was unaffected by any treatment (Fig. 7H), consistent with previous reports that phosphorylation at 14-3-3 binding sites does not impact Thr211 (Goransson et al, 2006). In HeLa cells, Ser396/400 phosphorylation was undetectable, and other MARK3 phosphopeptides showed lower phosphorylation and were unaffected by c604 (Fig. EV2C). These findings indicate that MARK3 regulation may differ across tumour types and suggest that Gwl controls MARK3 phosphorylation and activity in AML.

Among known MARK3 substrates (Peng et al, 1998; Yang et al, 2023), we identified ARHGEF2 Ser151 (Rho guanine nucleotide exchange factor 2 or GEF-H1) in our dataset, a site that disrupts its binding to microtubules (Sandi et al, 2017). The phosphorylation of ARHGEF2 at Ser151 and Ser163 increased upon mitotic arrest and decreased with c604 and CDK1i in P31/Fuj but not in HeLa cells (Fig. 7I, J). Aurora A kinase also phosphorylates ARHGEF2 at Ser886(Birkenfeld et al, 2007) in mitosis, but Aurora A activation during G2/M requires CDK1 activity (Van Horn et al, 2010), thus CDK1i should reduce ARHGEF2 Ser886 phosphorylation. As expected, Ser886 phosphorylation increased after mitotic arrest and was reduced by CDK1i but not by c604 in P31/Fuj or HeLa cells (Fig. 7I, J). Although c604 decreased phosphorylation of the CDK1 motif (Fig. 3D, E), this likely reflects altered mitotic progression rather than direct CDK1 inhibition, explaining its lack of effect on Ser886. Nevertheless, c604 reduced ARHGEF2 phosphorylation at Ser151, a MARK3-dependent site (Fig. 7I), indicating impaired MARK3 function and downstream signalling. This altered activity may disrupt microtubule dynamics, contributing to mitotic defects and reduced cell viability. Consistently, c604 treatment of primary AML cells tended to reduce the phosphorylation at three of the four detected MARK3 sites (Fig. 7K), although this did not reach statistical significance, likely reflecting patient heterogeneity. Together, these data support a role for Gwl in regulating MARK3 activity in AML cells.

## DISCUSSION

Gwl kinase is an emerging target for anticancer therapy in solid tumors (Rogers et al, 2018; Fatima et al, 2020; Zach et al, 2025; Castro & Lorca, 2018; Misra et al, 2023; Uppada et al, 2018; Vera et al, 2015). Here, we show that Gwl also plays a central role in regulating cell cycle progression and proliferation in AML cells. Strikingly, in AML, Gwl controls cell cycle progression via a noncanonical pathway. Indeed, Gwl pharmacological or genetic inhibition altered cell cycle progression, but, unlike in other models, these effects were not associated with reduced ENSA phosphorylation or diminished ENSA-B55α binding (Zach et al, 2025; Burgess et al, 2010; Hegarat et al, 2014). These findings were consistent across experimental systems and were also observed in primary AML cells, emphasising a leukaemia-specific regulatory mechanism distinct from other tumour types and models.

PP2A inactivation is a common feature in AML resulting from downregulation of PP2A-complex components or overexpression of endogenous inhibitors such as SET or CIP2A (Arriazu et al, 2016, 2020; Cristobal et al, 2012, 2011). Our data suggest that elevated basal phosphorylation of ENSA may contribute to PP2A suppression in AML, although the kinases mediating Gwl-independent ENSA phosphorylation remain to be identified.

Our findings suggest that Gwl regulates AML cell viability by modulating the phosphorylation of proteins involved in cytokinesis. We uncovered that Gwl phosphorylates MARK3 at residues involved in 14-3-3 binding. MARK3 controls microtubule dynamics and cell cycle progression (Machino et al, 2022; Peng et al, 1998), and is regulated by 14-3-3 protein binding and cellular localization (Goransson et al, 2006). MARK3 inhibition alters microtubule dynamics, and induces G2/M phase cell cycle arrest in glioma cells (Li et al, 2020), suggesting that MARK3 regulates cell cycle progression. Remarkably, c604 reduced MARK3 phosphorylation in primary AML cells and P31/Fuj cells. Interestingly, HeLa cells did not show this pattern of phosphorylation, which may suggest a different role of MARK3 across tumor types. The role of MARK3 in cellular signaling, particularly in cancer, appears highly context-dependent: in some settings it acts as a tumor suppressor, while in others it promotes tumor growth (Machino et al, 2022; Klingbeil et al, 2024; Mohseni et al, 2014; Owusu et al, 2019; Raut et al, 2025). Therefore, the differences observed between solid cancers and AML cells may also be related to the dual role of this kinase in tumor progression. Although its exact role in AML has not been fully elucidated, MARK3 is involved in the phosphorylation of MEF2C and chemoresistance in AML cells (Brown et al, 2018), suggesting that MARK3 plays a significant role in AML signaling.

MARK3 modulates the activity of ARHGEF2 (GEF-H1), an activator of RhoA signalling (Sandi et al, 2017) and regulator of microtubule dynamics (Sandi et al, 2017). ARHGEF2 is anchored to microtubules in an inactive state (Meiri et al, 2009, 2012), which is disrupted by MARK3 phosphorylation at Ser151 (Sandi et al, 2017). Notably, the increased phosphorylation of ARHGEF2 at Ser151 and Ser163 in mitotic P31/Fuj cells was reversed by c604 or CDK1i (Fig. 7I), indicating that Gwl inhibition modulates MARK3 activity. This effect was not observed in HeLa cells (Fig. 7J), suggesting a differential role of MARK3 in both cellular models. ARHGEF2 is a regulator of microtubule dynamics, controlling mitotic spindle orientation and cytoskeletal organization during cytokinesis (Bakal et al, 2005; Chang et al, 2008; Birkenfeld et al, 2007). Notably, ARHGEF2 controls mitotic spindle orientation in hematopoietic stem cells, and depletion of ARHGEF2 leads to an increase in abnormal divisions and alteration of hematopoiesis (Chan et al, 2021), suggesting ARHGEF2 is relevant for hematological cell division. Thus, if Gwl controls MARK3 activity during mitosis, c604 may impair ARHGEF2 activity, leading to improper microtubule dynamics and cytokinesis failure in AML cells. However, further studies are required to understand the role of ARHGEF2 in cell division, specifically in AML cells.

In summary, our study shows that in AML, Gwl kinase regulates mitotic progression through a mechanism independent of the canonical ENSA-PP2A-B55α pathway. This distinct signaling configuration in AML highlights the plasticity of cell cycle control in leukemic cells and expands our understanding of Gwl kinase function and mitotic regulation beyond the classical ENSA-PP2A-B55α axis.

## MATERIALS AND METHODS

### Cell lines and treatments

AML cell lines (Reagents and Tools Table) were maintained at a concentration of 0.5 × 106 cells/mL in RPMI 1640 medium supplemented with 10% FBS and 1% penicillin/streptomycin solution. OCI-AML3 cells were cultured in α-MEM supplemented with 10% FBS and 1% penicillin/streptomycin, and OCI-M1 cells were cultured in IMDM medium supplemented with 10% FBS and 1% penicillin/streptomycin. RPE1, HeLa, and HEK293T cells were cultured in DMEM supplemented with 10% fetal bovine serum (FBS) and 1% penicillin/streptomycin. Murine stromal cells, MS-5, were cultured in DMEM supplemented with 10% FBS and 1% penicillin/streptomycin. Cells were kept in an incubator at 37°C and 5% CO2, and regular mycoplasma testing was performed.

For inhibitor treatment, c604 (provided by Prof. Hochegger (Zach et al, 2025)), STLC (Selleckchem, Cat # S5522), nocodazole (Selleckchem, Cat # S2775), and RO-3306 (Selleckchem, Cat # S7747) were diluted in DMSO. The vehicle (DMSO) or the indicated concentration of the inhibitor was added.

Nocodazole or STLC arrest was induced by treating the cells for 18-20h in media containing 100 ng/mL nocodazole or 5 µM STLC. AML cells were collected by centrifugation, and RPE1 and HeLa cells were collected by mitotic shake-off and centrifugation, respectively. The cells were then resuspended in fresh media containing vehicle (DMSO) or the inhibitor at the required concentration and incubated for the required time. AML cells were seeded at a density of 0.25 × 106 cells/mL prior to cell cycle arrest or treatment with c604 or RO-3306.

### Cell viability, proliferation and cycle assays

AML cells were seeded in 96-well plates and treated with vehicle (DMSO) or 1-4 µM c604 for 72h, and viability was measured using a Guava PCA cell analyser. For cell cycle analysis, cells were fixed with EtOH 70%, stained with phosphorylated histone H3 Ser10 and PI/RNase solution. Samples were measured using a Guava PCA cell analyser (Guava Technologies Inc.), and data were analysed using CytoSoft (v2.5.7). Additional methodological details are provided as Supplementary Methods.

### Immunochemical methods

For immunofluorescence, cells were fixed with MetOH and stained against β-Actin and nuclei (using 4’,6-diamidino-2-phenylindole). Images were acquired using a Nikon Eclipse Ti-E inverted microscope equipped with a Yokogawa spinning disk and analysed using ImageJ/Fiji. To test the dynamics of ENSA-B55α complexes, B55α was immunoprecipitated using specific antibodies and the presence of ENSA in immunoprecipitates was measured by Western blot. For Western blots, proteins were separated by SDS-PAGE and transferred onto nitrocellulose membranes, which were incubated with the antibodies described in the main text, figure legends and Supplementary Methods.

### Generation of stable cell lines

Stable cell lines with IPTG inducible shRNA against Gwl were generated using lentiviral transduction as described in Supplementary methods.

### Phosphoproteomics analysis

Samples were digested and phosphopeptides were desalted and enriched with TiO2. Enriched phosphopeptides were analysed by label-free LC-MS/MS as previously described (Zach et al, 2025). Experiments were carried out in asynchronous or, when indicated, in prometaphase arrested cells using STLC (5 µM, 18-20 hours). Detailed procedures are described in Supplementary Methods.

### Kinase assays

To identify novel Gwl substrates, an in vitro kinase assay on fixed cells was performed as described by Al-Rawi et al. (Al-Rawi et al, 2023) using recombinant active Gwl kinases. To confirm direct MARK3 phosphorylation by Gwl, an in vitro kinase reaction using recombinant MARK3 and recombinant GST-Gwl kinases was carried out. Products of kinase reactions were identified and analyzed by LC-MS/MS as detailed in Supplementary Methods.

### Ethical Approvals

The patients provided informed consent for the storage and use of their blood cells for research purposes. The experiments were performed in accordance with the Local Research Ethics Committee as previously described (Casado et al, 2023, 2018).

### Culture and treatment of primary samples

Mononuclear cells from peripheral blood or bone marrow biopsies were isolated from tissue biobank facilities and stored in liquid nitrogen. Clinical details of these specimens are as in (Casado et al, 2023). These primary cells were cultured in media conditioned with MS5 bone marrow cells secretome, treated and processed for viability assays and phosphoproteomics-based LC-MS/MS as described above and in Supplementary Methods.

### Statistics

Statistical analysis was performed in R (v4.3.1) using base functions or ggpubr package. When not specified, statistical differences were evaluated using an unpaired Student’s t-test (p<0.05). Gene Ontology analysis of differentially expressed or phosphorylated genes/proteins was performed using the clusterProfiler package. The data were visualised using ggplot2. The ggseqlogo package was used for motif phosphorylation visualisation (Wagih, 2017). Metrics of response to c604 were calculated using PharmacoGx (Smirnov et al, 2016). Heatmaps were created using the ComplexHeatmap (Gu, 2022). For phosphoproteomics analysis, statistical differences were calculated using LIMMA (Ritchie et al, 2015), and then adjusted for multiple testing using the Benjamini-Hochberg procedure to control the false discovery rate (FDR). Differences were considered statistically significant when p<0.05 and FDR<0.15.

## DATA AVAILABILITY

The mass spectrometry proteomics data was deposited in the ProteomeXchange Consortium via the PRIDE (Perez-Riverol et al, 2025) partner repository with the dataset identifiers PXD068482 and 10.6019/PXD068482, and PXD069978 and 10.6019/PXD069978. Total proteomics and basal phosphoproteomics data from AML cell lines were obtained from Gerdes et al. (Gerdes et al, 2021) previously deposited with the identifiers PXD019591 and 10.6019/PXD019591. Genetic Dependency data were obtained from the DepMap database (https://depmap.org/portal): CRISPR (DepMap Public 25Q2) and RNAi (Achilles DEMETER2).

## AUTHOR CONTRIBUTIONS

S.M.M.G. designed and executed experiments and wrote the paper; S.M.M.G., T.S., V.R., P.R.C., R.Z., P.C, and N.A.N performed experiments; M.M. and W.R.F. contributed new reagents. I. I. B. and P.R.C. contributed new analytical tools; H.H. designed the experiments and revised the MS; P.R.C designed experiments and wrote the paper.

## COMPETING INTERESTS

The authors declare no competing interests.

## ACKNOWLEDGMENTS

We are thankful to Dr. Sam Wallis and the CRUK Barts Centre Microscopy Facility for their support with the image acquisition. We are thankful to the CRUK Barts Centre Mass Spectrometry Facility for its support. We thank the MRC PPU Reagents and Services Facility (MRC I PPU, College of Life Sciences, University of Dundee, Scotland, mrcppureagents.dundee.ac.uk) for the reagents and/or services indicated in this publication.

## FUNDING

This work was supported by funding from CRUK-C28206/A14499 (RZ, SMMG, MM, PRC) and CRUK-C15966/A24375 (PC, PRC). The Spanish Agencia Estatal de Investigación provided additional support (ATR2024-154531).

## Figure Legends

**Figure EV1. Effect of Gwl inhibition on AML cell proliferation.**

(A) Relative cell number of AML cells lines treated with different concentrations of Gwl inhibitor (c604) for 72h. Cell number is expressed as relative to DMSO treated cells. Values represent mean ± SD of 3 biological replicates. (B) Bar plots showing the response to c604 calculated as Area Above the Curve (AAC) from the data presented in A. Bar plots show the mean ± SD (n=3). (C) Gwl genetic dependency of cancer cell lines grouped by primary disease from RNAi Achilles Dataset. Statistical significance was calculated by unpaired t-Test between AML versus the rest of the cancer types (\**: p<0.05, : **p<0.01 and* : *** p<0.001). (D) Abundance of named proteins across the 14 AML cell lines tested and normalised signal intensity for the phosphorylation of ENSA/ARPP19 at Ser62 and Ser67 obtained from (Gerdes et al, 2021). (E) Distributions of cell cycle groups (2N, S, 4N, 4N+ and <2N) in AML cells treated with c604 (0.25 µM) for 24h and 48h of 3 independent biological replicates. Statistical significance was calculated by unpaired t-Test between treated cells versus non-treated (DMSO) groups (\**: p<0.05, **: p<0.01 and **\*\*\****: p<0.001). (F) Sequence motifs of significant decreased phosphopeptides in mitotic HeLa cells treated with c604.

**Figure EV2-MARK3 phosphorylation by Gwl.**

(A) Volcano plot shows phosphopeptides identified by LC-MS/MS in the kinase assay on fixed P31/Fuj cells. X-axis represents the FC (Log2 FC) of cells incubated with 8xHis-Gwl-2xFLAG versus cells incubated only with ATP (n=3). Blue and red colours indicate significant (p<0.05) phosphopeptides with decreased (FC< -0.7) and increased (FC> 0.7) phosphorylation compared to ATP only, respectively. Green dots show Gwl phosphopeptides. (B) Significant phosphopeptides identified in the kinase assay from A ranked by FC. Dashed pink line indicates the Log2 FC threshold of 0.7. (C) Box plot shows the normalised signal intensity for MARK3 phosphopeptides identified in P31/Fuj cells by LC-MS/MS. (D) Box plot shows the normalised signal intensity for MARK3 phosphopeptides identified in HeLa cells by LC-MS/MS. Unpaired t-Test was used to calculate the differences between the different treatments versus STLC treatment (mitotic) (\**: p<0.05 and \**\*: p<0.01).

## SUPPLEMENTARY INFORMATION

Supplementary Methods

References for supplementary methods

## Supplementary Methods

### Cell viability and proliferation assays

Cells were seeded in 96-well plates (10000 cells per well in 100 µL) and treated with vehicle (DMSO) or 1-4 µM C-604 for 72h. The final concentration of DMSO was kept at 0.2%. Cells were stained with Guava ViaCount Reagent (Cat. # 4000-0041) according to the manufacturer’s instructions. Samples were measured using a Guava PCA cell analyzer (Guava Technologies Inc.), and data were analyzed using CytoSoft (v2.5.7). At least three independent biological replicates were performed for each condition. Cell viability at the specified cell number was normalized to that of the control cells (DMSO-treated only).

### Cell cycle analysis

Cells were seeded in a T25 flask (0.25 × 10^6^ cells/mL) and treated with vehicle (DMSO) or C-604 for 24 and 48h. The final concentration of DMSO was kept at 0.2%. Cells were collected by centrifugation (500 g, 5 min, 4°C), washed with ice-cold phosphate-buffered saline (DPBS) containing phosphatase inhibitors (1 mM NaF and 1 mM Na_3_VO_4_) and fixed with cold 70% ethanol at -20°C for 12h. For double staining with phosphorylated histone H3 Ser10, EtOH was removed by centrifugation (1000 g, 10 min, 4°C), washed with DPBS, and stained with phosphohistone H3 Ser10 antibody in 1% BSA + DPBS + 0.1% Triton X-100 for 90 min at RT. Cells were then washed with 1% BSA + DPBS + 0.1% Triton X-100 (1000 g, 5 min, 4°C) and incubated with the secondary antibody in 1% BSA + PBS + 0.1% Triton X-100 for 90 min at RT. Finally, the cells were washed twice with DPBS (1000 g, 5 min, 4°C) and stained with PI/RNase solution (Guava® Cell Cycle Reagent, Cytek, Cat # SKU 4500-0220). Samples were measured using a Guava PCA cell analyzer (Guava Technologies Inc.), and data were analyzed using CytoSoft (v2.5.7). At least three independent biological replicates were performed for each condition.

For simple cell cycle staining, the cells were cultured, treated, and fixed as described above. After EtOH fixation and removal, the cells were washed in DPBS and stained with PI/RNase solution (BD Pharmingen™ PI/RNase Staining Buffer, Cat # 550825) for 30 min at RT. Samples were measured using a Guava PCA cell analyzer (Guava Technologies Inc.), and data were analyzed using CytoSoft (v2.5.7). At least three independent biological replicates were performed for each condition.

### Immunofluorescence

Cells were seeded in a T25 flask (0.25 × 10^6^ cells/mL) and treated with vehicle (DMSO) or C-604 for 48h. The final concentration of DMSO was kept at 0.2%. The cells were collected by centrifugation (500 g, 5 min, 4°C), washed with ice-cold DPBS, and fixed with cold 100% methanol at 4°C for 10 min. Methanol was removed by centrifugation (1000 g, 10 min, 4°C), and the cells were washed with DPBS. Cells were blocked with 3% BSA in DPBS + 0.1% Triton X-100 for 1 h at room temperature (RT). The cells were then incubated with primary antibodies in 3% BSA in DPBS + 0.1% Triton X-100 for 2 h at RT. Cells were washed with 1% BSA + DPBS + 0.1% Triton X-100 and incubated with secondary antibodies in 3% BSA in DPBS + 0.1% Triton X-100 for 2 h at RT. The cells were washed with 1% BSA + DPBS + 0.1% Triton X-100, followed by washing with DPBS. Nuclei were stained with DAPI solution, and the cells were washed with DPBS. Finally, the cells were resuspended in pure water. 5 µL drops of the cell suspension were spread onto Superfrost slides (VWR, Cat # 11836153001), dried, and mounted with Fluoroshield Mounting media (Abcam, Cat # ab104135). For all imaging experiments, cells were imaged using an Eclipse Ti-E inverted microscope (Nikon) equipped with a CSU-X1 Zyla 4.2 camera (Ti-E, Zyla; Andor), including a Yokogawa Spinning Disk, a precision motorized stage, and Nikon Perfect Focus, all controlled by NIS-Elements Software (Nikon). The Plan Apo VC 20x objective was used to acquire images. Images/projection images (from z-stacks) were subsequently generated and analyzed using ImageJ/Fiji (National Institute of Health, Bethesda, MD, USA). Only brightness and contrast were adjusted using the ImageJ/Fiji software. Blue color was changed to gray scale for presentation purposes only.

### Immunoprecipitations

Cells were washed with ice-cold DPBS supplemented with phosphatase inhibitors (1 mM NaF and 1 mM Na_3_VO_4_). The cells were then incubated with fresh 1 mM DSP (Thermo Fisher Scientific, Cat # A35393) reagent prepared in DPBS for 30 min at RT. Then, the DSP reaction was stopped by adding Tris-HCl pH 7.5 at a final concentration of 20 mM and incubating for 15 min at RT. The cells were then collected by centrifugation and washed with ice-cold DPBS containing phosphatase inhibitors. Next, the cells were lysed in IP buffer (50 mM Tris-HCl pH 7.4, 150 mM NaCl, 0.5% (v/v) Triton X-100, protease inhibitor cocktail containing EDTA, and phosphatase inhibitor cocktail PhosStop). Lysates were incubated at 4°C for 30 min. Following centrifugation at 16000 g for 15 min at 4°C, samples were incubated with the primary antibody (PP2A-B55-α Antibody) for 16h at 4°C. The antibodies were captured using Protein A/G PLUS-Agarose beads (Santa Cruz Biotechnology; 50% [v/v] in IP buffer) for 2h at 4°C and washed in TBS + 0.1% Triton X-100 supplemented with phosphatase inhibitors. Bound proteins were eluted and prepared for SDS-PAGE by incubation in SDS-PAGE sample buffer containing 50 mM dithiothreitol (DTT) and heating at 96°C for 5 min. The samples were separated using 4-12% Bis-Tris Protein Gels as described below.

### Western Blots

Cells were collected by centrifugation (500 g, 5 min, 4°C) and washed twice with icecold DPBS supplemented with phosphatase inhibitors (1 mM NaF and 1 mM Na_3_VO_4_). Cells were then lysed in RIPA buffer supplemented with 1 mM PMSF, 1x Protease Inhibitor Cocktail (Roche, Cat # 11836153001) and 1x PhosSTOP cocktail (Roche, Cat # 4906845001). Samples were sonicated for 10 cycles (30s ON 30s OFF) in a Diagenode Bioruptor® Plus, and insoluble material was removed by centrifugation (16000 g, 10 min, 4°C). The protein concentration was determined using a BCA Protein Assay Kit. Afterwards, the proteins were separated in 4-12% Bis-Tris Protein Gels (NuPAGE™ 4-12% Bis-Tris Protein Gels, ThermoFisher Scientific) and transferred onto nitrocellulose membranes. Blots were then blocked in 10% (w/v) skim milk (cat # 70166, Sigma) in TBS/T (Tris-buffered saline with 0.1% Tween-20) for 1h and incubated (4°C overnight) with a primary antibody solution (in 5% [w/v] skim milk in TBS/T, Reagents and Tools Table). Membranes containing the immunoprecipitated proteins were blocked with EasyBlocker 5% (GeneTex, Cat # GTX425858) in TBS/T for 1h at RT. Membranes were washed with TBS/T and incubated with the corresponding peroxidase-conjugated secondary antibody solution (5% [w/v] skim milk in TBS/T, Reagents and Tools Table) for 2h at RT. For the detection of phosphorylated proteins, blots were blocked in 5% BSA in TBS/T for 1h at RT, and then primary and secondary antibodies were incubated in 3% BSA in TBS/T. The membranes were stripped with 10 min incubation in stripping solution (ThermoFisher Scientific, Cat # 46430), followed by washing in TBS and blocking of the membrane. The antibody reaction was documented using the Amersham Imager 600RGB Chemidoc or BioRad ChemiDoc MP Imaging System using a chemiluminescence reagent (SuperSignal™ West Pico PLUS Chemiluminescent Substrate, ThermoFisher Scientific, Cat # 34080). Western blot quantification was performed using the Image Studio™ Lite Software (LI-COR Biosciences).

### Generation of stable cell lines using lentiviral transduction

Stable cell lines Lentiviral packaging was performed in HEK293T cells. Plasmids were transfected using a calcium/phosphate mammalian transfection kit (Promega, Madison, WI, USA). The virus was assembled from pCMV-VSV-G (Addgene, RRID:Addgene_8454), psPAX2 (Addgene, RRID:Addgene_12260) and packaged with plasmids containing the shRNA of interest (see below). Next, 300 μL of transfection mixture (HEPES-buffered saline, 2 μg plasmid DNA, 0.24 M CaCl_2_) was incubated at room temperature for 30 min at RT and added dropwise onto HEK293T cells (50-80% confluent). The transfected cells were incubated overnight at 37°C. The medium containing transfection reagents was replaced with fresh medium, and the cells were incubated for an additional 48h. The medium was collected, centrifuged, and filtered to retrieve viral particles. The infection medium was prepared by adding polybrene (5 μg/mL) and the filtered supernatant containing the virus to fresh complete medium. The infection medium was added to a cell culture of 0.5 × 10^6^ cells/mL. The cells were incubated for 72h. Cells were then collected by centrifugation, and selection was carried out for 10 days with 0.5 µg/mL of Puromycin dihydrochloride selection (Sigma, Cat # P8833-25MG). The cell lines generated were tested for mycoplasma regularly and grown in media supplemented with 0.5 µg/mL of Puromycin. Induction of shRNA was carried out by adding 1 mM IPTG to the medium prepared in water (Promega, Cat # V3955). The medium was replaced with fresh media containing 1mM IPTG every 48h. Cells without IPTG were grown under the same conditions as the control. The efficiency of shRNA knockdown was evaluated using western blotting. Plasmids containing shRNA: scramble shRNA pLV[shRNA]-LacI:T2A:PuroU6/2xLacO (Vector Builder, Cat # VB010000-9348prq); hMASTL[shRNA#1] pLV[shRNA]-LacI:T2A:PuroU6/2xLacO> (Vector Builder, Cat # VB240220-1161rgh).

### Sample preparation for phosphoproteomics analysis

Sample preparation for mass spectrometry analysis was performed as described previously (Zach *et al*, 2025). Cell pellets were lysed with a urea buffer (8 M urea in 20 mM HEPES, pH 8.0, supplemented with 1 mM Na_3_VO_4_, 1 mM NaF, 1 mM Na_2_H_2_P_2_O_7_ and 1 mM sodium β-glycerophosphate), sonicated for 30 cycles (30s ON 30s OFF) in a Diagenode Bioruptor® Plus, and insoluble material was removed by centrifugation (13000 rpm, 10 min, 4°C). The protein concentration was determined using the BCA protein assay kit. Extracted proteins (110 µg, 200 µL) were reduced with dithiothreitol (DTT, 12 mM) for 1h at 25°C and alkylated with iodoacetamide (IAM, 16.6 mM) for 30 min at 25°C. The samples were then diluted with 20 mM HEPES (pH 8.0) to a final concentration of 2 M urea and digested with equilibrated trypsin beads (50% slurry of TLCK-trypsin) overnight at 37°C. The beads were equilibrated by washing with 20 mM HEPES buffer (pH 8.0). After digestion and trypsin bead removal by centrifugation (2000 g, 5 min, 4°C), the peptide solutions were transferred into 96 Protein Lobind well plates and acidified by adding TFA to a final concentration of 1%. The samples were then desalted and subjected to phosphoenrichment using the AssayMAP Bravo platform (Agilent Technologies). For desalting, a peptide clean-up protocol (v3.0) was used.

Reverse phase S cartridges (Agilent, 5 μL bed volume) were primed with 250 μL of 99.9% acetonitrile (ACN) with 0.1% TFA and equilibrated with 250 μL of 0.1% TFA at a flow rate of 10 μL/min. The samples were loaded (770 μL) at 20 μL/min, followed by an internal cartridge wash with 250 μL 0.1% TFA at a flow rate of 10 μL/min. Peptides were then eluted with 55 μL of 70% ACN and 0.1% TFA in 96 protein LoBind plates containing 50 µL of 1M Glycolic Acid with 50% ACN and 5% TFA. Following the Phospho Enrichment v 2.1 protocol, phosphopeptides were enriched using 5µl Assay MAP TiO2 cartridges on the Assay MAP Bravo platform. The cartridges were primed with 100µl of 5% ammonia solution with 15% ACN at a flow rate of 300 μL/min and equilibrated with 50 μL of loading buffer (1 M glycolic acid with 80% ACN, 5% TFA) at 10 μL/min. Samples eluted from desalting were loaded onto the cartridge at 3 μL/min. The cartridges were washed with 50 μL of loading buffer and phosphopeptides were eluted with 25 μL of 5% ammonia solution with 15% ACN directly into 25 μL of 10% formic acid. Phosphopeptides were lyophilized in a vacuum concentrator and stored at - 80°C.

### Production and purification of recombinant active MASTL protein

Recombinant active 8xHis-MASTL-2xFLAG (His-MASTL-FLAG) was produced and purified at Sussex University by Prof. Helfrid Group using the Bac-to-Bac system (Anderson *et al*, 1995). Briefly, full-length MASTL (aa: 1-879) was cloned into the FastBac1 plasmid. Spodoptera frugiperda Sf9 cells were grown in 500 mL of Insect-Xpress medium (Lonza Bioscience) supplemented with penicillin and streptomycin. At a density of 2 × 106 cells, cells were infected with P3 virus at a Multiplicity of Infection (MOI) of 2. The cells were incubated for 48h at 27°C with 150rpm shaking. The cultures were then harvested by centrifugation at 1000 g for 20 min at 4°C. The harvested pellets were transferred into a Falcon tube and frozen at -20°C.

Cells were resuspended in gel filtration (GF) buffer (250 mM NaCl, 25 mM HEPES pH 7.5, 0.5 mM TCEP) supplemented with 10 mM imidazole and EDTA-free protease inhibitor tablets (Roche) on ice. Then, the cells were lysed on ice by sonication using a Vibra-Cell VCX 500 ultrasonic (amplitude of 35% with pulse sequences of 4 s ON, 12 s OFF, total time of 8 -10 min). The cell lysates were cleared by centrifugation at 37000 g for 1h at 4°C. Next, purification was performed using talon resin previously equilibrated with GF buffer containing 10 mM imidazole (wash buffer). The cell supernatant was incubated with Talon resin for 30 min at 4°C while rolling. The resin was collected by centrifugation at 700 g for 10 min, washed with wash buffer for 30 min while rolling, and centrifuged again (700 g, 10 min). This step is repeated several times. Following the washing steps, the resin was loaded back into a gravity column, and the protein was eluted from the resin with elution buffer (GF buffer with 250 mM imidazole). After elution from the talon resin, the protein sample was incubated with Flag resin pre-equilibrated with GF buffer for 30 min at 4°C with rolling. The two wash steps were repeated with GF buffer. Next, to elute the protein, the column was incubated for 15 min with GF buffer supplemented with 100 μg/mL 3x Flag peptide. This was repeated to 2-3 times. Finally, the protein was pooled, concentrated using a Vivaspin column, and loaded onto a preequilibrated size-exclusion column (Cytiva Superdex Superdex 200 Increase 5/150).

### *In vitro* kinase assay on fixed cells

An *in vitro* kinase assay on fixed cells was performed as described by Al-Rawi et al (Al-Rawi *et al*, 2023). 8 × 10^6^ cells P31/Fuj cells were fixed with 1% PFA supplemented with phosphatase inhibitors (1x PhoSTOP, Roche) for 10 min at RT, and then permeabilized with 100% ice-cold methanol for 10 min on ice. After washing the cells in ice-cold DPBS supplemented with phosphatase inhibitors (1x PhoSTOP, Roche), reaction tubes were prepared by blocking 2 × 10^6^ cells in blocking buffer (40 mM Tris-HCl pH7.5 supplemented with 5% BSA and 1x PhosSTOP) for 10 min at RT. Cells were then washed by centrifugation at 15000 g for 30s and placed in 400 µL of phosphorylation master mix (40 mM Tris pH 7.5, 0.1 mg/ml BSA, 1 mM ATP, 10 mM MgCl_2_, 1mM DTT and 1x PhosSTOP). Then, recombinant active MASTL kinases, GST-MASTL (0.5 µg, SinoBiological, Cat # M72-10G) or His-MASTL-Flag (1 µg, provided by Prof. Helfrid) were added and incubated for 90 min at 25°C. As a negative control, cells with only the phosphorylation master mix and no kinase were incubated. The reactions were quenched by sequential washes with quenching buffer (40 mM Tris pH 7.5, 0.1 mg/ml BSA, 11 mM EDTA, and 1x PhosSTOP), 13000 g for 30s at 4°C. Finally, cells were lysed in digestion buffer (100mM triethyl ammonium bicarbonate [TEAB] pH 8.5, 2 mM MgCl_2_ and 5 U Benzonase) for 30 min at 37°C. Protein was then digested by adding 1.25 µg Trypsin protease for 16 h at 37°C followed by another addition of 1.25 µg trypsin for 4h. The samples were acidified by the addition of trifluoroacetic acid (TFA) at a final concentration of 1%. Peptide desalting was performed using C18 + GC carbon tips (TopTip with Filter C18 + Carbon 10-200 µL, cat # TF2MC18.96, Glygen). C18 tips were equilibrated by adding 200 µL of 100% ACN twice (1500 g, 2 min, 4°C) and then washed twice with 200 µL of 99% H_2_O/1% ACN/0.1% TFA (1500 g, 2 min, 4°C). The samples were then loaded into the tips by centrifugations at 1500 g 2 min at 4°C. After sample loading, the tips were washed twice with 200 µL of 99% H_2_O/1% ACN/0.1% TFA (1500 g, 2 min, 4°C). Finally, peptides were eluted into new 2mL Lo-Bind tubes by adding 100 µL of 1 M glycolic acid in 50% ACN and 5% trifluoroacetic acid (TFA). This step is repeated twice. The sample volumes were normalized to 300 µL with 1 M Glycolic acid/ 80% ACN/ 5% TFA and incubated for 5 min with TiO2 (50 µL per sample of 500 µg/ml of TiO_2_ in 1% TFA). Samples were then loaded into empty spin tips by centrifugation for 30s at 1500 g. Samples were sequentially washed by centrifugation with 100 µL of 1 M Glycolic acid/ 80% ACN/ 5% TFA, then with 100 µL of 100 mM ammonium acetate in 25% ACN) and three times with 100 µL of (90% H_2_O/10% ACN). Phosphopeptides were eluted four times with 50 µL of 5% NH_4_OH and 10% ACN into new 1.5mL Lo-Bind Eppendorf tubes. Finally, the eluents were snap-frozen in dry ice for 15 min, dried in a speed vac overnight, and phosphopeptide pellets were stored at -80°C.

To check protein phosphorylation by western blotting, after the kinase reaction and quenching, cells were resuspended in 70 µL of cell extraction buffer (1mM HEPES, 10 mM EDTA, 2% SDS, 1x COMPLETE protease inhibitor cocktail, and 1x PhosSTOP). The extracts were then sonicated for 30 min (30s ON, 30s OFF), and crosslinking was reversed by heating the samples at 95°C for 50 min. Proteins in the lysates were then reduced by adding 30 mM DTT and mixed with sample buffer for loading. Proteins were separated using SDS-PAGE, as described above.

### *In vitro* kinase assay for LC-MS/MS analysis

To confirm MARK3 phosphorylation by MASTL, an *in vitro* kinase reaction using recombinant MARK3 (MRC PPU Reagents and Services facility, College of Life Sciences, University of Dundee, Scotland, Cat #DU1296) and recombinant GST-MASTL (SinoBiological, Cat # M72-10G) kinases was carried out as described below. Five micrograms of 6XHis-MARK3 were incubated in phosphorylation master mix reaction (40 mM Tris pH 7.5, 50 µM ATP, 10 mM MgCl_2_, 1mM DTT and 1x PhosSTOP) in the presence or absence of GST-MASTL (0.25 µg) for 30 min at 37°C. The reaction was stopped by the addition of 8 M Urea, and the samples were reduced with dithiothreitol (DTT, 12 mM) for 1h at 25°C and alkylated with iodoacetamide (IAM, 16.6 mM) for 30 min at 25°C. The samples were then diluted with 20 mM HEPES (pH 8.0) to a final concentration of 2 M urea and digested with equilibrated trypsin beads (50% slurry of TLCK-trypsin) overnight at 37°C. The beads were equilibrated by washing with 20 mM HEPES buffer (pH 8.0). After digestion and trypsin bead removal by centrifugation (2000 g, 5 min, 4°C), peptides were desalted using C18 + GC carbon tips, and phosphopeptides were enriched with TiO2 as indicated for the kinase assay on fixed cells.

### LC-MS/MS Analysis

The phosphopeptides were re-suspended in 8 µL of reconstitution buffer and sonicated for 1 min at RT. After centrifugation (13000 rpm, 10 min, 4°C), 5μL was loaded onto an LC-MS/MS system. This consisted of a nano flow ultra-high-pressure liquid chromatography system (UltiMate 3000 RSLC nano (Dionex) coupled to a Q Exactive Plus using an EASY-Spray system. The LC system used mobile phases A (3% ACN: 0.1% FA) and B (100% ACN: 0.1% FA). The peptides were loaded onto a μ-pre-column and separated on an analytical column. The gradient was as follows: 1% B for 5 min and 1% B to 35% B over 90 min, after which the column was washed with 85% B for 7 min and equilibrated with 3% B for 7 min at a flow rate of 0.25 µL/min. Peptides were nebulized into an online connected Q-Exactive Plus system operating with a 2.1s duty cycle. Acquisition of full scan survey spectra (m/z 375-1,500) with a 70,000 FWHM resolution was followed by data-dependent acquisition, in which the 15 most intense ions were selected for HCD (higher energy collisional dissociation) and MS/MS scanning (200-2,000 m/z) with a resolution of 17,500 FWHM. A 30s dynamic exclusion period was enabled with an exclusion list with a 10ppm mass window. The overall duty cycle generated chromatographic peaks of approximately 30s at the base, which allowed the construction of extracted ion chromatograms (XICs) with at least ten data points.

### Phosphopeptides identification and quantification

Peptide identification from MS data was automated using a Mascot Daemon workflow in which Mascot Distiller generated peak list files (MGF) from RAW data, and the Mascot search engine matched the MS/MS data stored in the MGF files to peptides using the SwissProt Database restricted to Homo sapiens (SwissProt_2024_03. fasta). Searches had an FDR of ∼1% and allowed two trypsin missed cleavages, mass tolerance of ±10 ppm for the MS scans and ±25 mmu for the MS/MS scans, carbamidomethyl Cys as a fixed modification and oxidation of Met, PyroGlu on N-terminal Gln, and phosphorylation of Ser, Thr, and Tyr as variable modifications. A percolator was then applied to improve the discrimination between correct and incorrect spectrum identifications (Brosch *et al*, 2009). Pescal was used for label-free quantification of the identified peptides as described previously (Wilkes *et al*, 2017). The software constructed XICs for all peptides identified in at least one LC-MS/MS run across all samples. The XIC mass and retention time windows were 7 ppm and 2 min, respectively. Peptides were quantified by measuring the area under the XIC peaks. Individual peptide intensity values in each sample were normalized to the sum of the intensity values of all the peptides quantified in that sample. Finally, phosphoproteomics data were processed and analyzed using a public bioinformatics pipeline developed in an R environment (https://github.com/CutillasLab/protools2/). The normalized data were centered, and log2 scaled and 0 values were replaced by the minimum feature value minus one. Statistical differences (p-values) were calculated using LIMMA (Ritchie *et al*, 2015), and then adjusted for FDR using the Benjamini-Hochberg procedure. Differences were considered statistically significant when p-values were <0.05 and FDR<0.15.

### Culture and treatment of primary samples

For the collection of primary culture media (conditioned media), MS-5 cells were plated in T125 flasks at an initial density of 1×10^6^ cells in complete DMEM. Cells were then incubated at 37°C overnight, the growth medium was removed and replaced with H5100 medium (STEMCELL Technologies, Cat # 05150) containing 1% P/S, and cells were incubated at 37°C for 48h. Then, the growth medium was collected and centrifuged at 500 g for 5 min. Then, the medium was mixed 1:1 with fresh H5100 medium containing 1% P/S and stored at 4°C until use.

Mononuclear cells from peripheral blood or bone marrow biopsies were isolated from BCI or FIMM tissue bank facilities and were stored in liquid nitrogen. Cells were thawed at 37°C, transferred to 50 mL falcon tubes, and incubated for 5 min at 37°C with 500 µL of DNAse Solution (Sigma Aldrich, Cat# D4513-1VL; resuspended in 10 mL of PBS). Then, 10 mL of 2% FBS in PBS was added, and the cell suspension was centrifuged at 525 g for 5 min at RT. The supernatant was discarded, and the cells were resuspended in 10 mL of conditioned media supplemented with human interleukin-3 (PeproTech, Cat # 200-03), human granulocyte colony-stimulating factor (PeproTech, Cat # 300-23), and human thrombopoietin (PeproTech, Cat # 300-18), at a final concentration of 20 ng/mL. To evaluate the effect of C-604 on the viability of primary AML cells, primary cells were seeded in 96-well plates (30000 cells per well in 100 µL) and treated with vehicle (DMSO) or 1-4 µM C-604 (C-604) for 72h. Viability was assessed using the Guava ViaCount Reagent and Guava PCA cell analyzer (Guava Technologies Inc.), as indicated for AML cell lines.

For phosphorproteomic analysis, primary cells were thawed as described above, seeded in a T25 flask (8 × 10^6^ cells), and incubated for 3 h in an incubator at 37°C and 5% CO_2_. The cells were then treated with vehicle (DMSO) or C-604 and incubated for 2h. The final concentration of DMSO was kept at 0.2%. For cell harvesting, cell suspensions were centrifuged for 5 min at 525 g at 4°C, and the pellets were washed twice with DPBS supplemented with phosphatase inhibitors (1 mM Na_3_VO_4_ and 1 mM NaF). Pellets were transferred to low-protein binding tubes and stored at -80°C. The phosphoproteomic analysis was performed as described above.

## Structured Methods - Reagents and Tools Table

*Instructions: Please complete the relevant fields below, adding rows as needed. The following page provides an example of a completed table and additional instruction for entering your data in the table*.

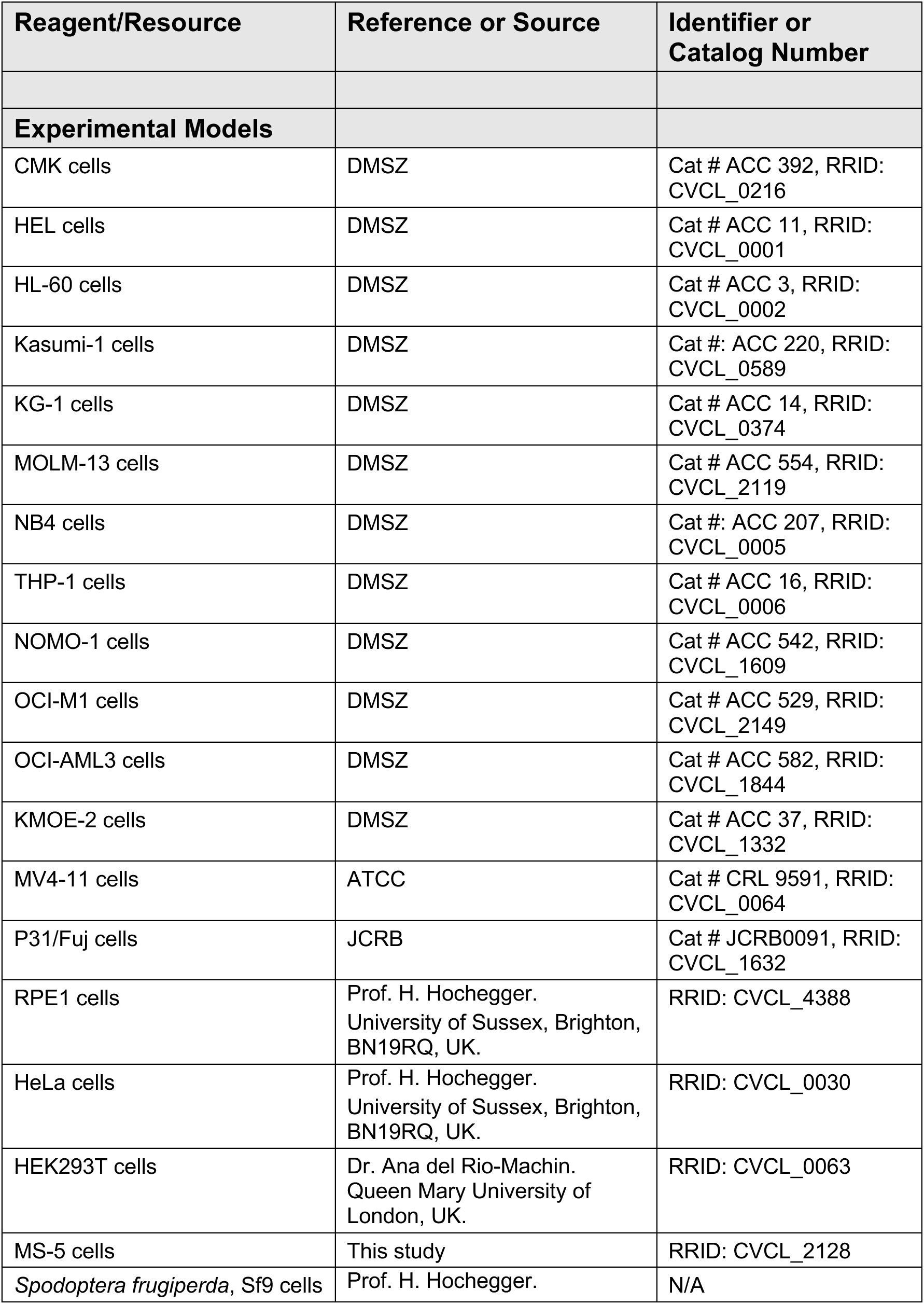
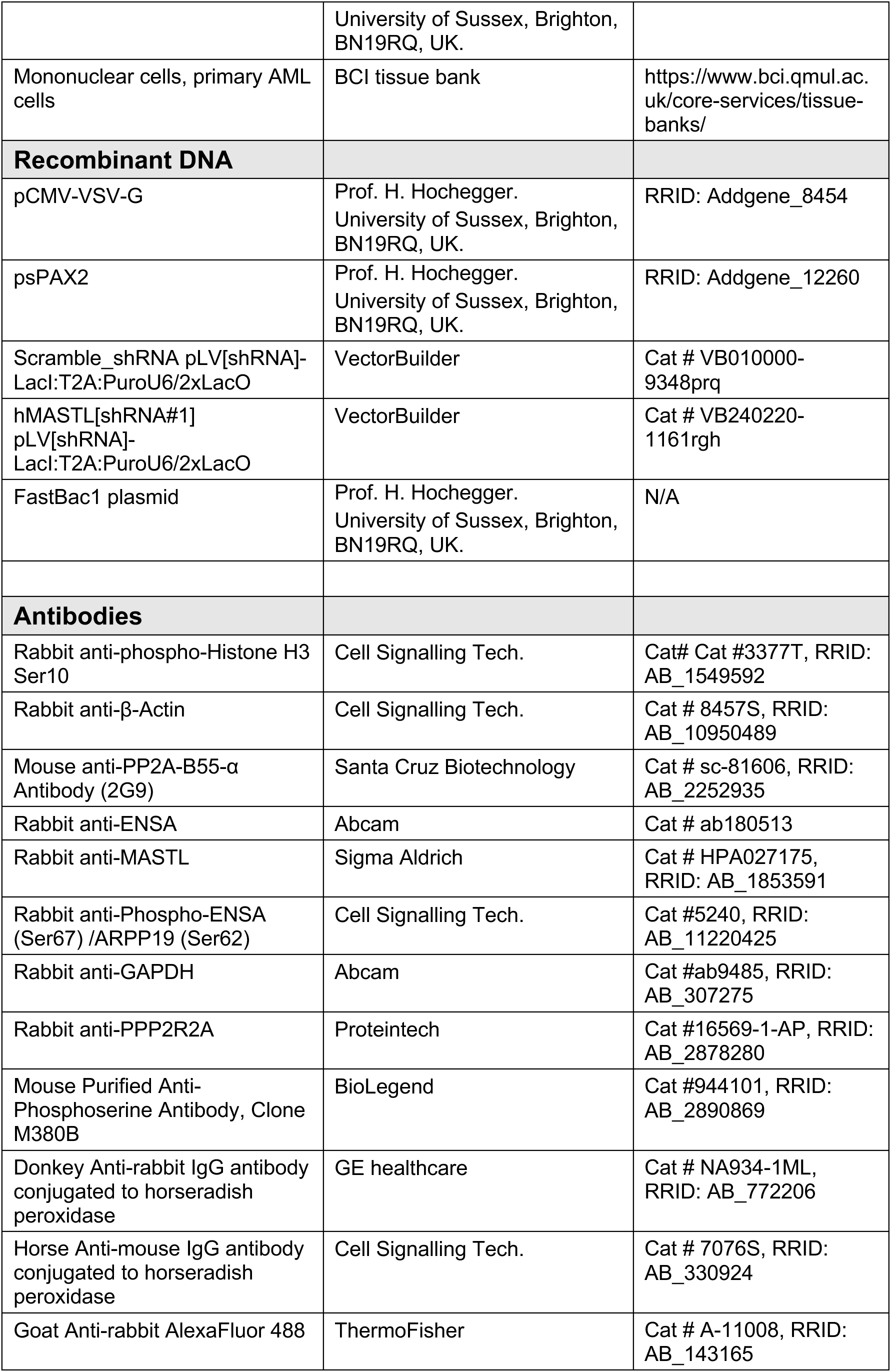
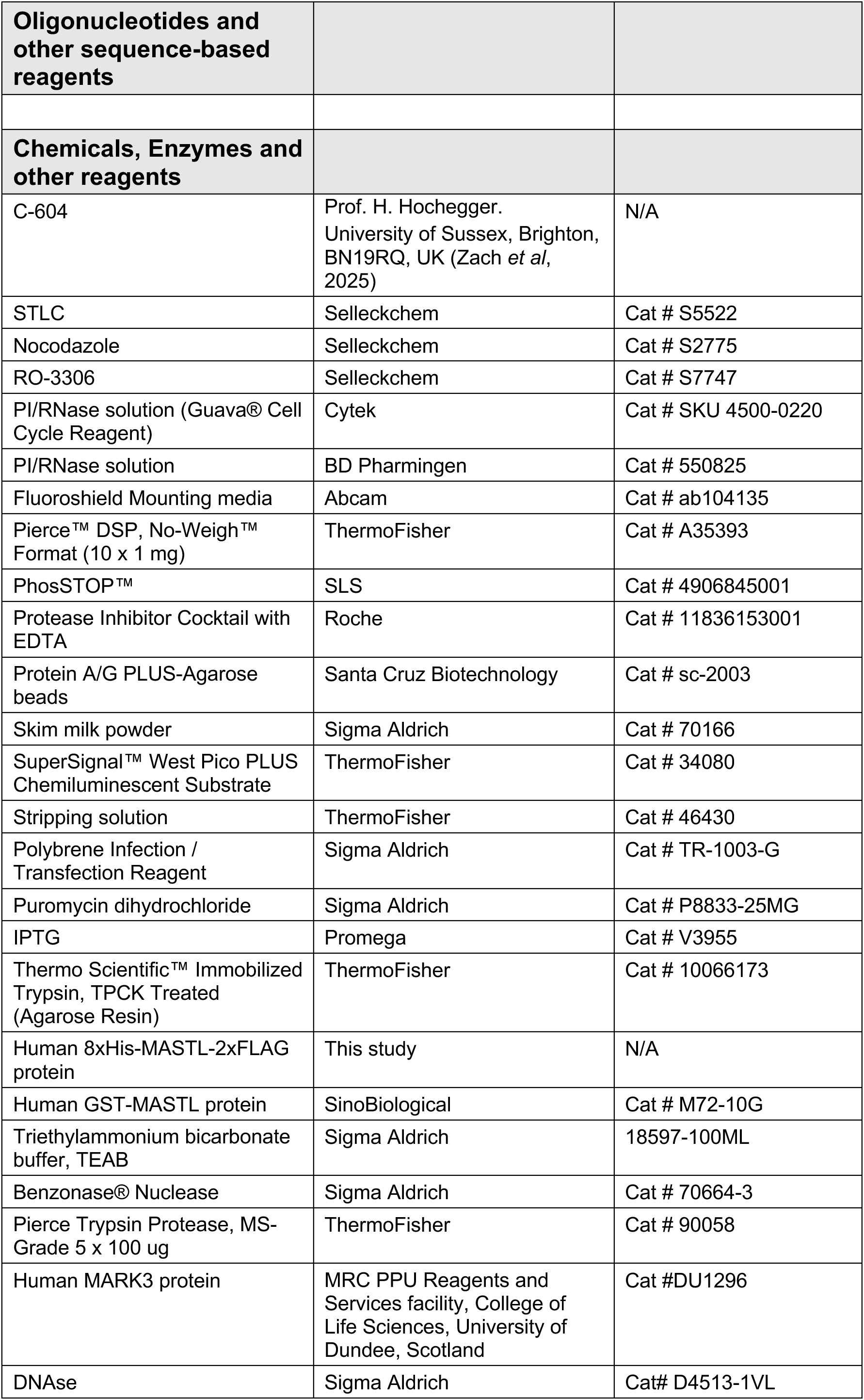
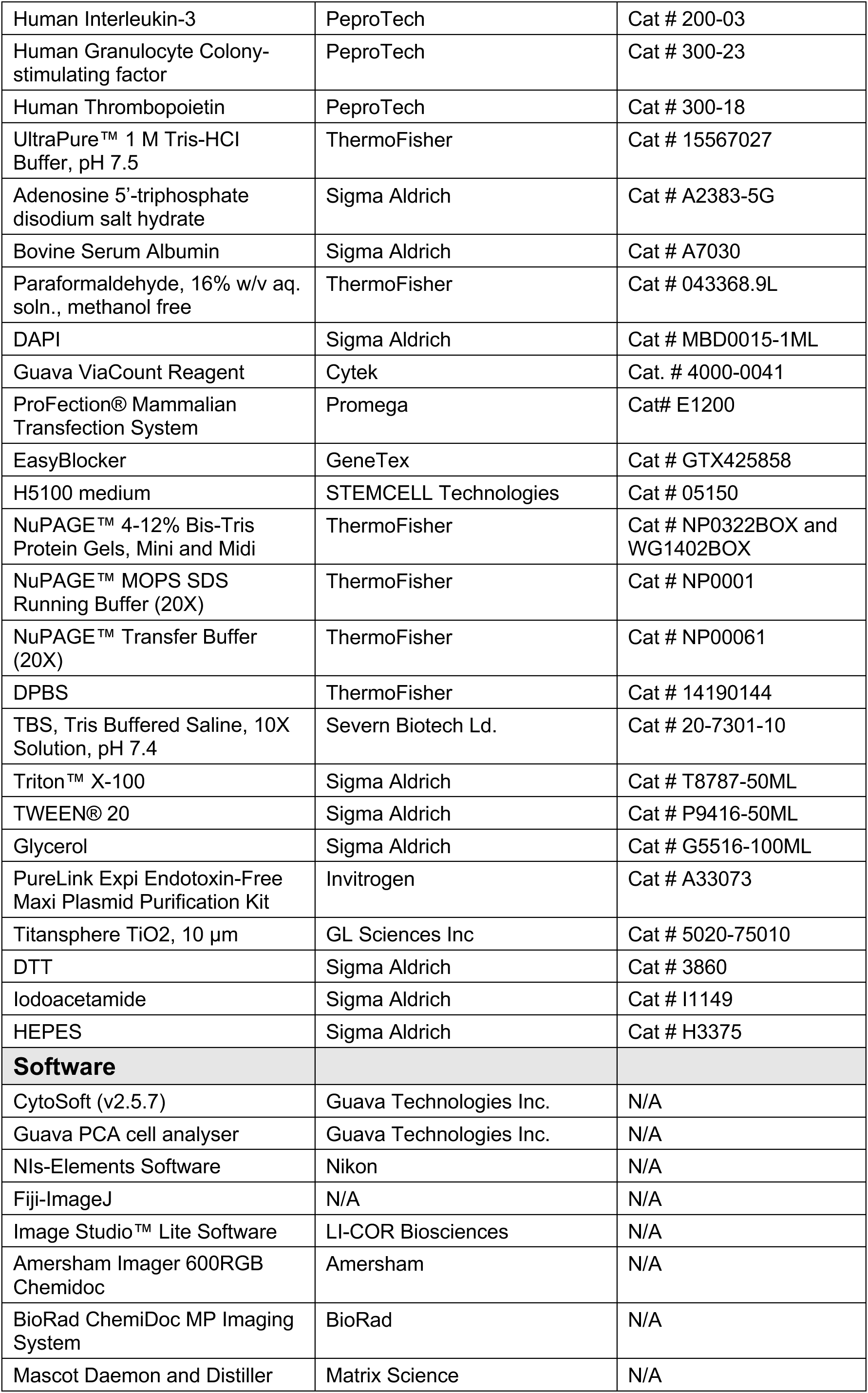
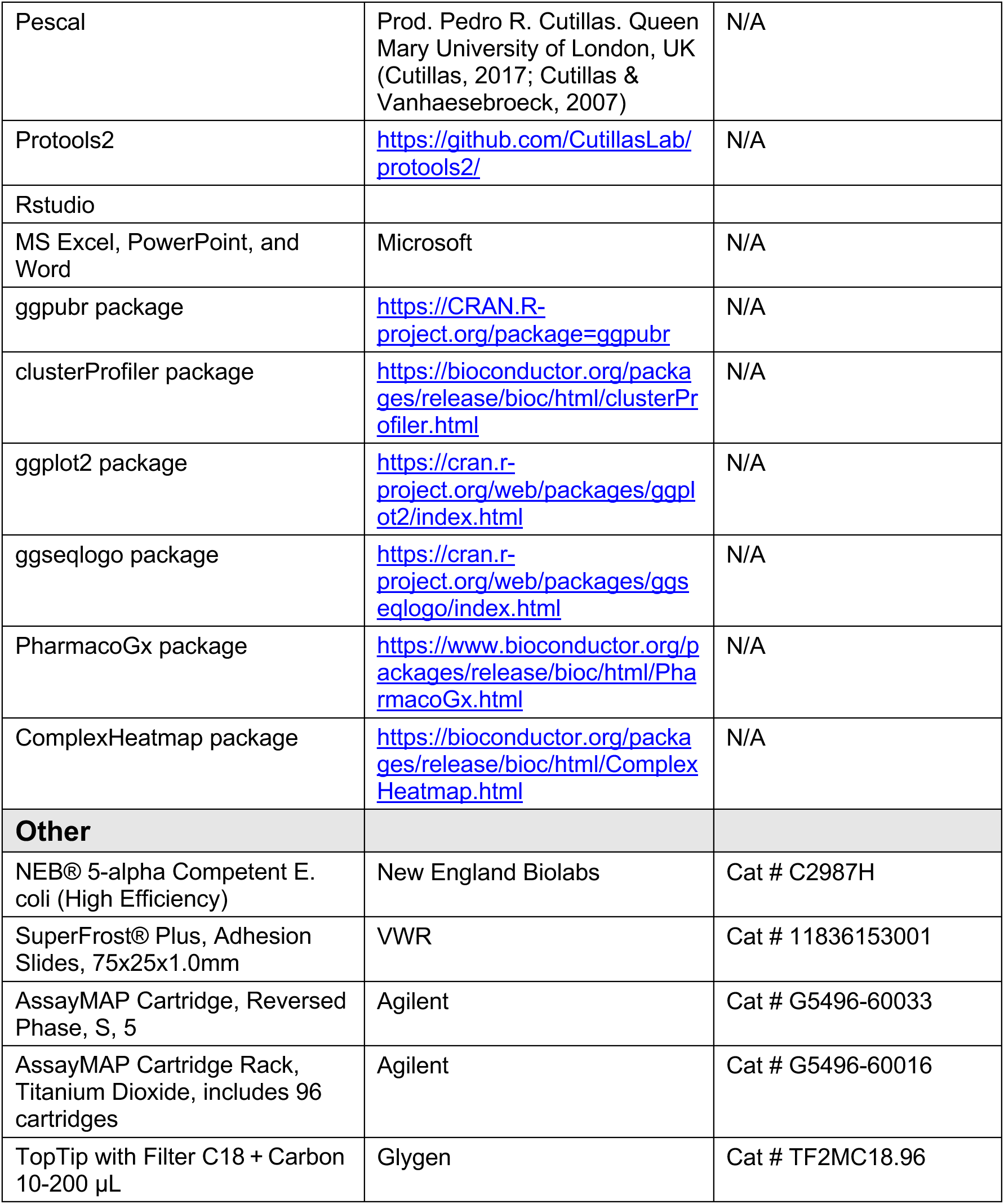

